# Footwear-specific biomechanical and energetic responses to 8 weeks of training in advanced footwear technology

**DOI:** 10.1101/2024.11.20.624520

**Authors:** Justin R. Matties, Joshua D. Kerr, K. Michael Rowley

## Abstract

**Background:** While the acute effects of advanced footwear technology (AFT) on running biomechanics and efficiency have been extensively studied, the longitudinal effects of AFT use are unknown.

**Objective:** The purpose of this study was to investigate the effects of using advanced footwear technology (AFT) versus traditional racing flats during running workouts on relative running economy (RE) and energetic cost (EC), and explore associations between changes in footwear-specific biomechanics and performance.

**Methods:** Thirteen competitive runners were randomly assigned Nike Vaporfly Next% 3 (VP) or Nike Rival Waffle 5 (FL) for an 8-week intervention and completed pre- (PRE) and post-intervention (POST) lab testing in both shoes. Weekly training data, including mileage, workouts, and soreness, were collected via questionnaires. Subjects ran two 3-min trials at a self-selected cross-country race pace, with sagittal plane ankle and metatarsophalangeal (MTP) joint kinematics and kinetics recorded and analysed. Subsequently, four 5-min trials were completed at a self-reported submaximal pace, in a randomized order, and shoe-specific RE and EC were calculated and reported as “VP% benefit” – the percentage RE or EC improvement in VP versus FL. Pearson’s correlations were tested between RE outcomes and exploratory biomechanics measures, and independent samples t-tests tested for group differences in shoe-specific and overall efficiency changes from PRE to POST.

**Results:** VP% benefit increased from PRE to POST in VP trained runners and decreased in FL trained runners. VP trained runners decreased ankle plantarflexion velocity in VP, increased ankle dorsiflexion velocity in VP, and decreased MTP plantarflexion velocity in FL. Correlations revealed several significant associations between metabolic outcome measures, and biomechanical and training variables.

**Conclusions:** Metabolic data support a footwear specificity of training principle (habituation), where training in VP may enhance a runner’s ability to benefit from VP on race day. Associations reveal that this may be due to changes in footwear-specific biomechanics at the ankle and MTP joints. Future research should explore the potential mechanisms through which runners adapt to AFT to maximize their performance benefits.

**Key Points:**

- The acute metabolic benefits and reduced mechanical demands in advanced footwear technology (AFT) have previously been demonstrated in distance runners. In an 8-week footwear intervention study, runners who trained in AFT increased their metabolic benefit from AFT and runners who trained in traditional racing flats decreased acute AFT metabolic benefits.
- Changes in ankle and metatarsophalangeal joint mechanics partially explain these changes in AFT metabolic benefits.
- Data from this research support the findings of an earlier pilot study and replicate a habituation effect with AFT use during hard effort workouts, suggesting runners may consider using AFT prior to competition to maximize performance benefits.

## 1 Introduction

Advanced footwear technology (AFT) has contributed to significant improvements in elite distance running performances from the 5k to marathon, with multiple long-standing world records being broken with the use of AFT [1–5]. The acute performance benefits of AFT have been measured in laboratory settings with 2.5-4% improvements in metabolic efficiency (ME); with ME expressed as running economy (RE), the amount of oxygen (ml·kg^-1^·min^-1^) used, or as energetic cost (EC), the amount of energy used (W·kg^-1^) used, to run at a given speed [6–8]. Currently the improvements in ME and changes in biomechanics in AFT are attributed to: (1) an embedded carbon fibre plate or similar longitudinally rigid structure; (2) a high stack height of supercritical midsole foam that is lightweight, compliant, and resilient; and (3) a more pronounced rocker geometry [9–11]. In contrast to AFT, traditional racing flats typically have a lower stack height and use non-supercritical midsole foams that are less compliant, or “cushioned” [6, 8]. Outside of the lab, retrospective analyses of elite race performances have found significant (1-3%) improvements in elite race times since the introduction of AFT, with these improvements increasing with longer distance races [1, 2, 4, 5].

These acute improvements in performance observed in AFT have been attributed to biomechanical changes observed in AFT compared to more traditional racing footwear. Some of these acute biomechanical responses to AFT include decreased ankle positive and negative work, decreased ankle plantarflexion velocities that likely result in more efficient triceps surae contractions, decreased metatarsophalangeal (MTP) joint work, and decreased energy loss at the MTP joint [7, 12, 13]. Which of these changes in mechanics in AFT are primarily contributing to the observed 2.5-4% improvement in ME from AFT is still unclear. Acute decreases in post-workout muscular soreness have also been noted with AFT use.

Despite the growing popularity of using AFT in running workouts, there is a lack of knowledge about the effects of training in these shoes longitudinally. Previous minimalist footwear interventions have observed increases in foot and plantarflexor strength [14, 15], which align with the acute effects of minimalist footwear on the ankle and metatarsophalangeal (MTP) joint mechanics [16]. Therefore, it is possible that the acute effects of AFT, including decreased metabolic expenditure, decreased mechanical demand on the ankle and MTP joints, and reduced muscular soreness [17], may stunt positive training adaptations in runners training in AFT. Alternatively, these acute effects of AFT, including reductions in post-workout muscular soreness [17], may allow for increased training loads.

With the assumption that runners will opt to race in AFT to realize the acute performance gains, the impact of training footwear on their ability to benefit from AFT on race day is an important consideration for competitive and elite runners. If an adaptation period exists for AFT, runners may habituate to a specific shoe through (1) changes in movement coordination after repeated exposure to the footwear condition [18], or (2) through footwear-specific musculoskeletal adaptations that may improve gait efficiency as suggested by minimalist footwear interventions [14, 15, 19–21]. With no clear definition for these hypothesized phenomena in the literature, we refer to a more recent definition of habituation as the biomechanical, neural, sensory, and cognitive changes that occur after regular, repeated exposure [22–24].

A previous pilot 8-week intervention was conducted a year prior to the present intervention [25]. Both the pilot and present interventions were 8 weeks in length, an intervention length based on previous minimalist footwear interventions who observed changes between footwear intervention groups [15, 18–21]. Results from the pilot study suggested a potential specificity of training principle with footwear, or habituation effect, where runners’ footwear-specific ME improved in their intervention shoe. Runners who completed the pilot intervention in traditional racing flats also improved their average ME, independent of testing shoe, to a greater extent over the 8 weeks, though with only 2 runners completing the intervention in this group we have little confidence in this finding. With a small, unbalanced sample of 8 runners, these findings need to be replicated and serve as the hypotheses of the present study.

The purpose of this study is to investigate the effects of using AFT versus traditional racing flats during running workouts on overall ME and VP% benefit, and to explore biomechanical trends that are associated with changes in footwear-specific performance. Comparisons are made between training groups to assess differences in how ME, VP% benefit, and footwear-specific biomechanics change over the intervention. Based on pilot data from Matties et al. [25], we hypothesize that: (1) AFT trained runners will increase their acute AFT ME benefits and improve overall ME and (2) traditional flat-trained runners will decrease acute AFT ME benefits and improve overall ME to a greater extent than AFT-trained runners.

## 2 Methods

### 2.1 Participants

Thirteen competitive cross-country runners (male: n = 10; female: n = 3; age: 22.40 ± 3.71 years; height: 176.40 ± 9.85cm; body mass: 67.63 ± 11.07kg) were recruited from men’s and women’s National Collegiate Athletic Association (NCAA) Division I, Division II, community college, and club cross-country teams in the California Bay Area and completed the intervention (Figure 1, Table 1). Participants were eligible if they were competing in at least one cross-country race in the fall 2023 season, were between 18 and 35 years of age, and regularly ran at least 30 miles per week. Exclusion criteria included any lower extremity musculoskeletal problem that required more than a one week break from running in the previous three months, or any serious injury or surgery on the lower extremities. All participants provided informed consent to participate, and all procedures were approved by an Institutional Review Board (CSUEB-IRB-2022-150).

**Fig. 1.**
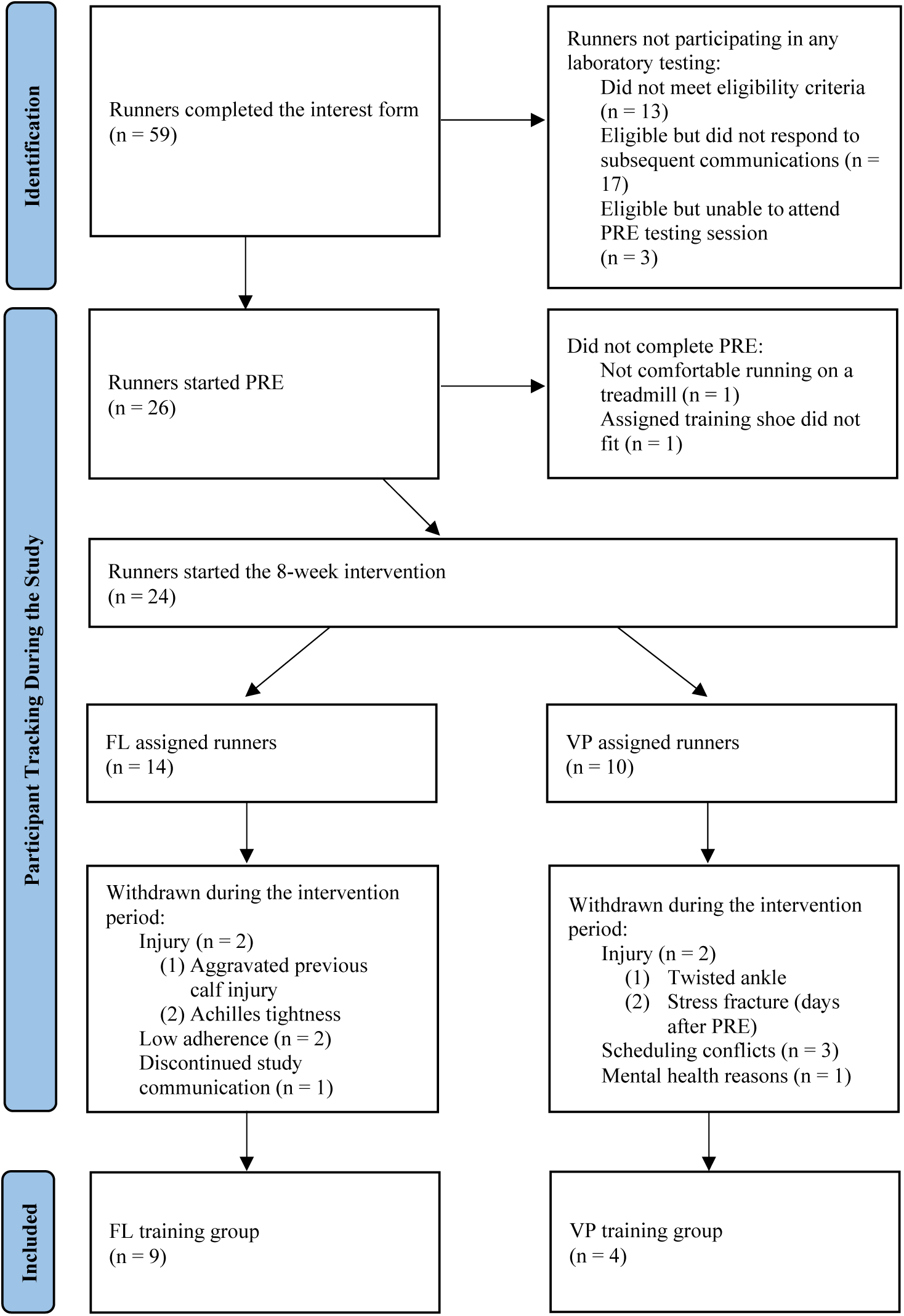
Participant selection and adherence flowchart.

**Table 1.** Participant characteristics table with means and standard deviations (SD) shown. Traditional racing flat (FL) and Vaporfly (VP) training groups are separated.

|  | FL Training Group (Mean<br>± SD) | VP Training Group<br>(Mean ± SD) |
| --- | --- | --- |
| n | 9 | 4 |
| Male/Female | 6/3 | 4/0 |
| Age (years) | 22.33 ± 4.27 | 22.50 ± 2.52 |
| Height (cm) | 174.11 ± 10.55 | 181.58 ± 6.28 |
| Mass (kg) | 63.32 ± 9.65 | 77.32 ± 7.76 |
| Self-Selected Race Pace<br>(km·hr<sup>-1</sup>) | 17.01 ± 1.80 | 17.93 ± 0.96 |
| Self-Selected Submaximal Pace<br>(km·hr<sup>-1</sup>) | 14.71 ± 1.42 | 15.23 ± 0.86 |
| Self-Selected Race Pace (m·s<sup>-1</sup>) | 4.72 ± 0.50 | 4.98 ± 0.27 |
| Self-Selected Submaximal Pace<br>(m·s<sup>-1</sup>) | 4.08 ± 0.39 | 4.23 ± 0.24 |
| Weekly Mileage (miles) | 49.52 ± 16.50 | 50.51 ± 7.35 |
| Weekly Workout Mileage<br>(miles) | 8.75 ± 2.55 | 7.92 ± 1.20 |
| Adherence to Shoe Type | 87.39 ± 12.10% | 97.92 ± 3.61% |
| Intervention Length (Weeks) | 8.46 ± 1.18 | 8.86 ± 0.46 |

### 2.2 Footwear

The Nike ZoomX Vaporfly Next%3 (VP) (Figure 2a) served as the AFT model. The Nike Zoom Rival Waffle 5 (FL) (Figure 2b) was used as a traditional racing flat model. Both VP and FL were used in laboratory testing and provided to participants as training intervention footwear based on group assignment. One pair of VP and FL in a shoe size range from US men’s size 5 to size 12 were instrumented and set aside for PRE and POST testing. No laboratory testing shoes accumulated more than 15km.

**Fig. 2.**
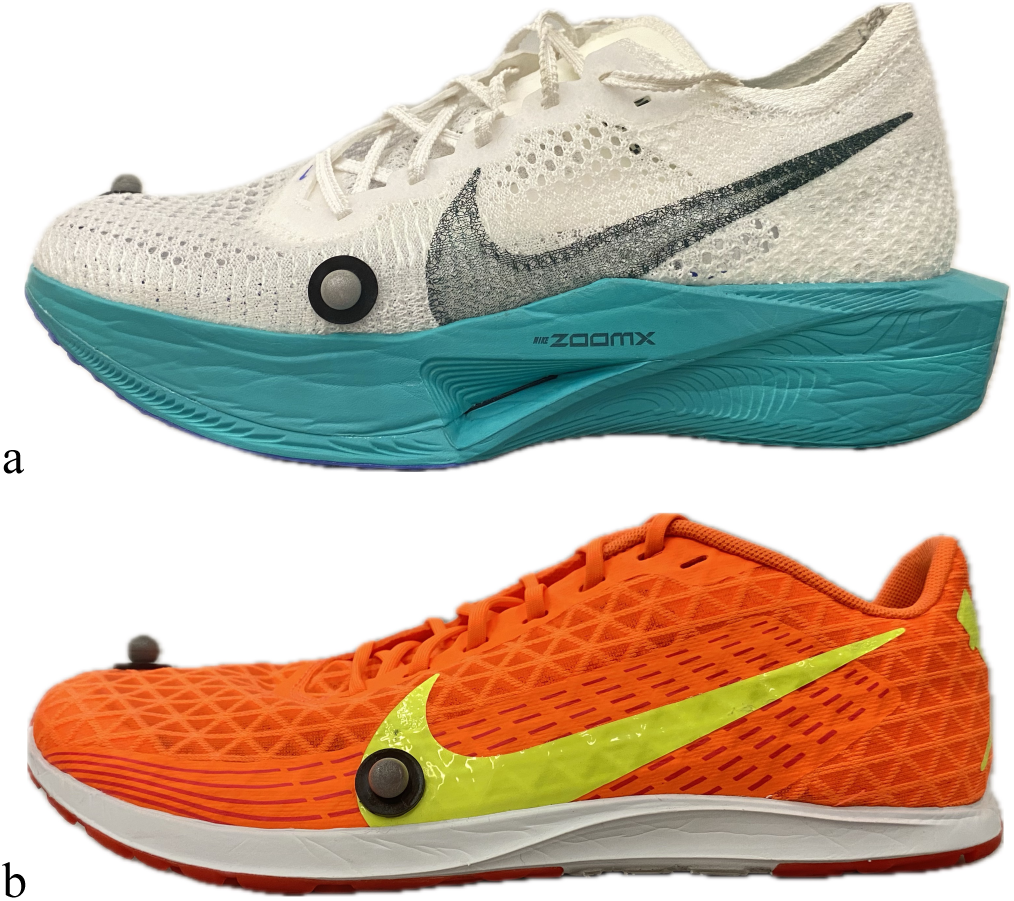
(a) Nike ZoomX Vaporfly Next% 3 (VP); (b) Nike Zoom Rival Waffle 5 (FL)

### 2.3 Instrumentation

Participants completed identical pre- (PRE) and post- (POST) intervention lab testing in both VP and FL. During PRE, participants gave informed consent; completed an intake running questionnaire [26] that captured details about training status, load, injury history, and running shoe preference [27]; and testing shoes were fitted. The intake questionnaire also asked for runners to detail previous training shoe and AFT experience. The questionnaire showed that six of the thirteen runners that completed the intervention had experience with at least one commercially available AFT model, though models were from different brands (Nike: n = 5; Saucony: n = 3; Hoka: n = 2; ASICS: n = 1; Adidas: n = 1). With the variation of previous AFT used both within and between runners, and only one runner using a Nike Vaporfly Next% 3 three months prior to the study, prior AFT history was not included in group stratification. In PRE and POST, participant height and weight were recorded.

Participants were instrumented with a modified Cleveland Clinic 6DoF retroreflective lower-body marker set and ran on a force plate (1000Hz, AMTI, Watertown, MA, USA) instrumented treadmill (Treadmetrix, Park City, UT, USA). Markers were placed bilaterally on the ASIS, PSIS, and iliac crest of the pelvis. Foot markers were glued to the shoe upper laterally to the 1st and 5th MTP joints, as well as superior to the distal phalanx of the first digit. These markers were used to distinguish the toe segment and acquire movement data of the MTP joints between the toe segment and the rest of the foot [28]. Plastic plates with four non-colinear markers were used to track the thigh and shank. Plastic plates with superior, inferior, and lateral heel markers were secured to the heel of each shoe. Calibration only markers were placed on the greater trochanters, medial and lateral femoral condyles, and medial and lateral malleoli. All markers were placed by the same individual who was trained in marker placement, with a second trained individual ensuring proper placement. Motion data were captured with an 8-camera Vicon Nexus System (100Hz, Vicon Motion Systems LTD, United Kingdom).

### 2.4 Procedures

First a Hans Rudolph metabolic mask and a Garmin HRM Pro heart rate monitor (Garmin HRM-Pro, Garmin Ltd, Olathe, KS, USA) were fitted to participants. Metabolic data were collected using a mixing chamber with 30 second averages reported (TrueOne 2400, Parvo Medics, Salt Lake City, UT, USA). Participants were then given a safety briefing and an explanation of the Borg’s 6 to 20 Rating of Perceived Exertion (RPE) scale [29, 30]. Participants then ran a warm-up in their own shoes for a self-selected duration between 15 and 20 minutes on the treadmill at a self-selected pace (same pace and time repeated for POST). Participants wore the metabolic mask for the final 5 minutes of the warmup for familiarization during PRE and POST.

Once warmed up, participants removed their shoes and were led through the single leg hopping protocol to measure vertical stiffness (Kvert) [31, 32]. Participants completed 20 seconds of single leg hopping at 2.2Hz (MetroTimer, ONYX Apps, Bellevue, WA, USA) on their right leg, followed by the left leg [31, 33, 34]. Hopping trials were conducted on the same stiff, instrumented treadmill as running trials (Treadmetrix, Park City, UT, USA) with the belt secured in place. Forces were measured by AMTI force plates (1000Hz, AMTI, Watertown, MA, USA).

After the completion of hopping trials, participants ran two 3-minute trials [7, 35] at self-selected goal cross-country race pace (10k pace for male runners: 17.95 ± 1.05 km⋅hr^-1^; 6k pace for female runners: 15.10 ± 1.12 km⋅hr^-1^) with 7-10 minutes rest to change shoes and secure shoe markers. During these race pace trials, biomechanical data were collected in each shoe (VP/FL) and slow-motion video was recorded in the sagittal plane and level with the treadmill surface at 240Hz (iPhone 11 Pro Max, Apple Inc, Cupertino, CA, USA). Three-dimensional kinematics and kinetics were recorded during minute 2 of these trials [35]. Subjects did not wear a metabolic mask during race pace trials. Following the race pace trials, retroreflective markers were removed. Then subjects completed four 5-minute trials with 5 minutes rest in between each [6, 8, 36, 37] at a self-selected submaximal pace (male runners: 15.34 ± 0.97 km⋅hr^-1^; female runners: 13.29 ± 0.68 km⋅hr^-1^) [8, 36, 38] while wearing a face mask. Trial paces were kept constant in PRE/POST. Shoe order during testing followed an ABBAAB pattern for all six trials [6, 8, 26, 37]. Shoe A was randomly selected for each participant in PRE and kept the same for POST.

### 2.5 Intervention

Participants were randomly assigned to either the VP or FL training group for an 8-week intervention in which they were instructed to complete their hard running workouts and races in their assigned shoe. Hard running workouts were defined as threshold/tempo runs, track workouts, longer interval sessions, or fartlek runs that typically occur twice a week in collegiate cross country training programs. Easy, recovery, medium-long, long, and easy “double” runs were not considered to be hard running workouts. Runs that were not considered hard running workouts were done in their standard training shoe of choice. Participants were asked to maintain at least an 80% adherence rate in their assigned shoe during hard workouts and races through the 8-week intervention. Runners were advised to use their assigned intervention shoe for 25-50% of their first intervention workout and 50-75% of their second intervention workout.

Participants completed a weekly questionnaire to record weekly mileage, workout/race details, shoe usage, perceived exertion, and muscular soreness. Perceived muscular soreness was recorded for the day of and the day following a workout/race in the participant’s assigned shoe using a five-point Likert scale. RPE for each intervention workout/race was assessed using a 10-point scale [29, 30, 39]. Participants were also asked to detail any injuries sustained during the intervention period.

### 2.6 Signal Processing

Metabolic data were collected using a mixing chamber analysis. Gross oxygen consumption (VO_2_) and respiratory exchange ratio (RER) data were averaged for the last 2 minutes of the submaximal trials [6–8, 37, 40]. Averaged VO_2_ data was used to calculate VP% benefit (RE) and overall ME improvement (RE) (ml·kg^-1^·min^-1^) for all subjects. VO_2_ and RER data were used to calculate VP% benefit (EC) and overall ME improvement (EC) (W·kg^-1^) using non-protein-based RER equations for participants that completed all trials with an RER<1 [41]. VP% benefit was calculated in terms of RE and EC using Equations 1 and 2 respectively. Overall ME improvements were calculated as the percent difference between the average RE and EC across all four submaximal trials in PRE and POST, with a positive percent improvement value representing greater efficiency, or decreased average EC or VO_2_ uptake.

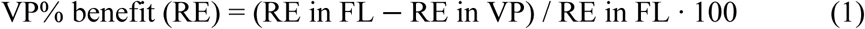

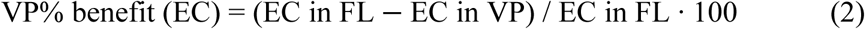

Biomechanics were captured during the race pace trials only. Marker trajectories and force plate data during race pace trials were low-pass filtered with a 20Hz Butterworth filter [42–44]. After processing the kinematic data, the latest 10 strides with clean data (e.g. no segments missing, no thigh or shank marker cluster slipping) were exported to Visual 3D for analysis (Visual 3D, C-Motion Inc., Germantown, MD, USA). A 50N vertical ground reaction force (GRF) threshold was used to determine gait events (footstrike/toe-off) and these were checked visually in every trial. Visual inspection of slow-motion video was used to determine footstrike technique (FST) in VP and FL during PRE and POST. Operational definitions of FST included: rearfoot strike (RFS) - striking with only the posterior third of the foot; forefoot strike (FFS) - striking with only the anterior half of the foot; and midfoot strike (MFS) was used to classify anything in between RFS and FFS [45–47]. Changes in main outcome measures within subsets of subjects based on FST will be explored. Visual 3D was used to calculate sagittal plane hip, knee, ankle, and MTP joint powers, moments, angles, and angular velocities bilaterally using a six degree of freedom model. Joint angles are prone to small differences in marker placement between testing sessions and between shoe conditions, therefore joint velocities were calculated using the derivative of the respective joint angle during stance phase.

To narrow the scope of this analysis, only ankle and MTP joint mechanics were considered due to their more immediate interaction with the shoe and the relatively greater effects of AFT on these joint mechanics relative to the knee and hip [7, 12]. Additionally due to the complexities of the current study design, a difference between shoes (DBS) metric was calculated for biomechanical variables during PRE, and another DBS was calculated during POST. DBS changes from PRE to POST (ΔDBS) were found for each biomechanical variable of interest using Equation 3 below.

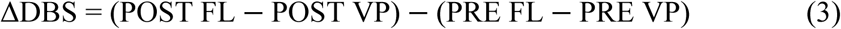

Kvert (kN·m^-1^) was calculated as the ratio of the maximal vertical ground reaction force (vGRF) and the lowest point of vertical centre of mass (COM) displacement using a double integration technique assuming the COM position is zero at take-off, referred to as SP1 and described in detail by Hébert-Losier and Eriksson [34]. Analysis was done using a custom Matlab script (Matlab R_2023a, The MathWorks Inc., Natick, MA, USA). All take-off and landing event detections were confirmed visually using the raw and filtered GRF, and individual hops were excluded from analysis if automatic event detection was not correct or hop duration was not within 5% of the prescribed 2.2Hz. Included hops were then rank ordered by stiffness with the middle 22 included in the average for each leg. Kvert values reported are an average of left and right legs.

### 2.7 Statistical Analysis

Separate correlational analyses using Pearson correlation coefficients were performed to describe associations between outcome measures and the two main metabolic outcome measures including overall RE improvement and changes in VP% benefit (RE). Subject-based (mass, Kvert) and intervention-based variables (weekly mileage, workout mileage, soreness, and RPE) were used in correlational analyses with overall RE improvement. PRE to POST changes in subject-based and biomechanical variables were used in correlational analyses with changes in VP% benefit (ΔVP% benefit). Associations were tested between ΔDBS and changes in ΔVP% benefit. Correlations are reported with p-values corrected for multiple comparisons. Significant correlations (p < 0.05) are further described to compare differences between FL and VP trained runners. Analyses were conducted in R (R Studio, PBC, Boston, MA, USA).

Effect sizes were determined for differences between shoe intervention groups from PRE to POST. Due to the small, uneven group sizes in the present study, Hedges’ g was used as a conservative descriptive statistic [48]. Significance levels calculated with an independent samples Student’s *t*-test assuming equal variances are used to compare main outcomes of interest, including VP% benefit, overall ME improvements, Kvert, and muscular soreness differences from PRE to POST between training intervention groups. P-values less than 0.05 were considered significant. All trends are described even when non-significant to contribute novel information to the literature and provide future directions of inquiry and hypothesis building for more rigorous testing in the future.

## 3 Results

### 3.1 Participation

#### 3.1.1 Intervention and Adherence

Intervention lengths were similar between the FL and VP training groups (FL: 8.46 ± 1.18; VP: 8.68 ± 0.46 weeks; p = 0.3208). Average total weekly mileage during the intervention was similar between training groups (FL: 49.52 ± 16.50; VP: 50.51 ± 7.35 miles per week; p = 0.4909). The average amount of weekly workout mileage, different from the total weekly mileage in that it only includes the distance run during hard workouts and races, was similar between groups though slightly higher in the FL training group (FL: 8.75 ± 2.55; VP: 7.92 ± 1.20 miles; p = 0.1979).

Of the 24 runners that completed PRE and started the training intervention, 13 finished the intervention and completed POST (Figure 1). Of the 13 runners who completed the training intervention, 9 runners were assigned to the FL training group and 4 were assigned to the VP training group. The FL training group’s adherence rate on average was 87.38 ± 12.1%, with two runners dropping below the requested 80% adherence rate, with individual adherence rates of 70% and 69%. These runners are included in analyses due to the small sample size of this study and their avoidance of carbon-plated AFT during intervention workouts/races. These adherence rates include the first week of the intervention where FL trained runners were strongly encouraged to slowly introduce FL into their training workouts/races. All runners in the VP training group exceeded the 80% requested adherence rate with an average adherence of 97.92 ± 3.61%, which was significantly higher than the FL training group (p = 0.0427).

#### 3.1.2 Soreness and RPE

On average, ratings of muscular soreness, where a rating of 1 represented no soreness and a rating of 5 represented extreme soreness, were non-significantly higher in FL trained runners compared to VP trained runners during the remainder of the day following a workout/race (FL: 2.76 ± 0.60; VP: 2.3 ± 0.59; p = 0.1347; T = 1.3751; DF = 10; g = 0.2440) (Figure 3a). Differences between training groups were less pronounced the day after a workout/race (FL: 2.39 ± 0.57; VP: 2.19 ± 0.69; p = 0.3351; T = 0.4387; DF = 10; g = 0.1033) (Figure 3b). Participants’ average RPE for workouts and races during the intervention were similar, with VP trained runners reporting slightly, though non significantly, lower RPE’s than the FL trained runners (FL: 6.69 ± 0.88; VP: 6.18 ± 0.84; p = 0.1429; T = 0.8933; DF = 10; ES (Hedge’s g) = 0.1836) (Figure 3c). Here an RPE of 0 represented rest and a rating of 10 represented maximal exertion.

**Fig. 3.**
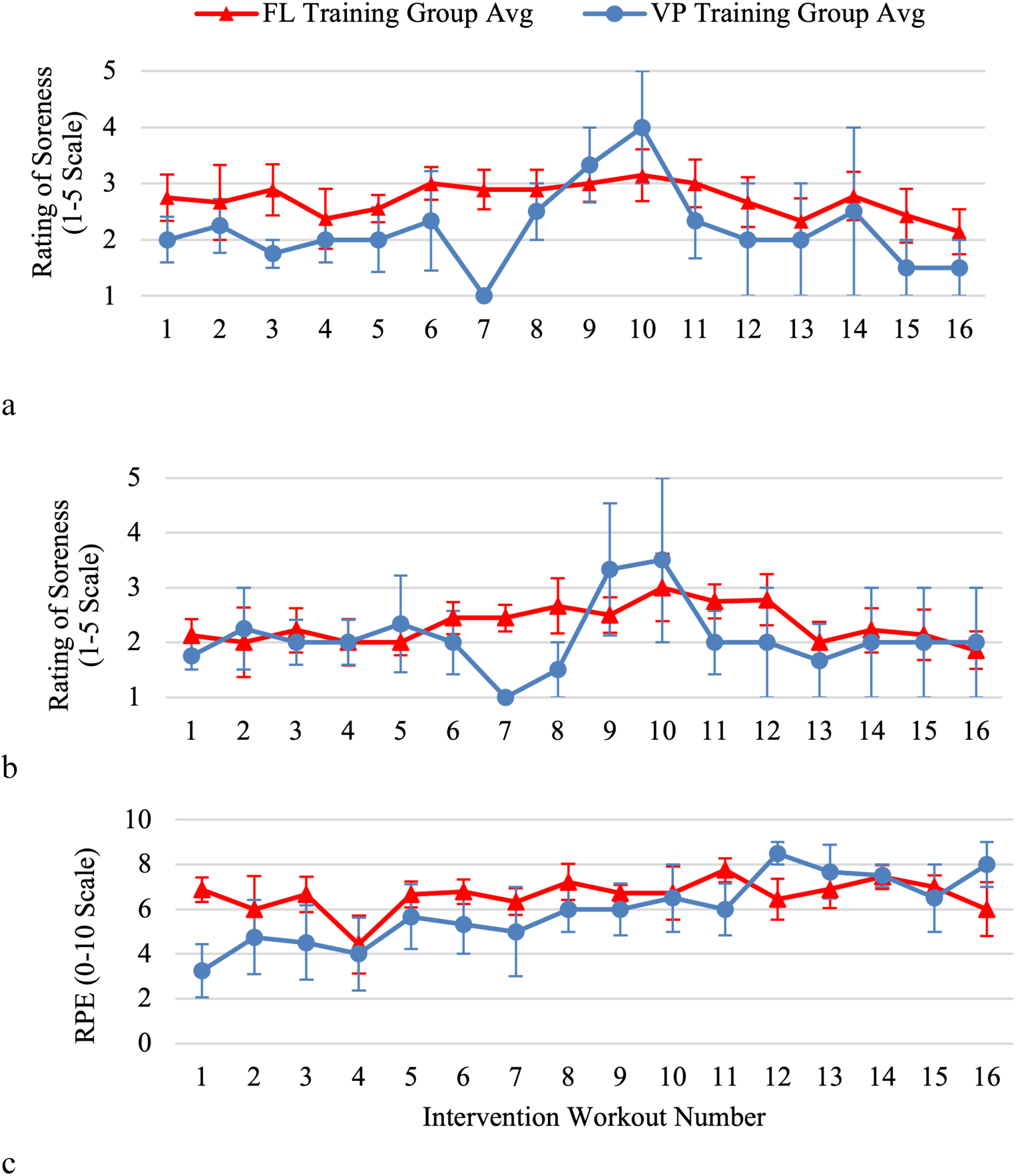
Perceived muscular soreness and rating of perceived exertion (RPE) graphs for the 8-week training intervention. Standard error bars are shown for each rating of soreness and RPE. Average perceived muscular soreness within each intervention training group is noted for both the remainder of the day after intervention workouts (a), and the following day (b). (c) Average Borg’s RPE (0-10 scale) within each intervention training group is given for each intervention workout

### 3.2 Overall ME and Functional Test Changes

#### 3.2.1 FL and VP Training Group Comparison

Comparing FL and VP intervention groups, there were no significant differences in overall ME improvement between FL trained runners (RE: 1.68 ± 4.76%; EC: 2.43 ± 3.30%) and VP trained runners (RE: 1.32 ± 4.31%; EC: 0.95 ± 5.17%) (RE: p = 0.4357, g = 0.0288, T = 0.1656, DF =11; EC: p = 0.3291, g = 0.1363, T = 0.4593, DF = 8) (Table 2). Two FL and one VP intervention group subjects exceeded an RER of 1.0 in at least one submaximal trial. These runners’ RE data is included in the present RE analysis but excluded from the EC data analysis. Kvert increased in FL trained runners (PRE: 26.58 ± 3.14 kN·m^-1^; POST: 27.94 ± 4.44 kN·m^-1^; ΔKvert: 1.36 ± 3.85 kN·m^-1^) and decreased in VP trained runners (PRE: 31.85 ± 3.91 kN·m^-1^; POST: 31.44 ± 3.59 kN·m^-1^; ΔKvert: −0.41 ± 1.02 kN·m^-1^), though group differences in ΔKvert were non-significant (p = 0.1577, g = 0.2014, T = 1.0711, DF = 8).

**Table 2.** Overall running economy (RE) and energetic cost (EC) values (mean ± standard deviation) for Vaporfly (VP) and traditional racing flat (FL) training groups during pre- (PRE) and post-testing (POST). The percent improvement from PRE to POST is also shown, with a lower RE and EC value in POST equating to a positive percent improvement, signifying improved efficiency. P-values represent between group differences in the change from PRE to POST (% Improvement)

|  | FL Training Group |  |  | VP Training Group |  |  | p |
| --- | --- | --- | --- | --- | --- | --- | --- |
|  | PRE | POST | % Improvement | PRE | POST | % Improvement |  |
| Running Economy ($\text{VO}_2$; $\text{ml}\cdot\text{kg}^{-1}\cdot\text{min}^{-1}$) | $45.47 \pm 6.03$ | $44.64 \pm 5.72$ | $1.68 \pm 4.76$ | $48.27 \pm 2.58$ | $47.67 \pm 3.14$ | $1.23 \pm 4.31$ | 0.4357 |
| Energetic Cost ($\text{W}\cdot\text{kg}^{-1}$) | $16.26 \pm 1.94$ | $15.89 \pm 2.19$ | $2.43 \pm 3.3$ | $16.75 \pm 0.19$ | $16.6 \pm 1.05$ | $0.95 \pm 5.17$ | 0.3291 |

#### 3.2.2 Associations with Overall ME Changes

Of the 7 subject-based and intervention-based variables tested in a correlational analysis with overall RE change, change in runner mass was the only variable that had a statistically significant association with changes in overall RE in all runners (r = 0.5667; p = 0.0434), though this is likely because the RE metric is normalized to subject mass. Moderate, though non-significant, associations were also observed between overall RE improvement and weekly workout mileage (r = −0.4828; p = 0.1119) and workout day soreness (r = 0.4012; p = 0.1961) (Table 3). When exploring associations among all variables included, three additional statistically significant associations were observed: (1) increases in Kvert and increases in mass were significantly positively associated (r = 0.8892; p = 0.0006); (2) workout day soreness and day after workout soreness were significantly positively associated (r = 0.8443; p = 0.0006); and (3) day after workout soreness and weekly mileage were significantly negatively associated (r = −0.7051; p = 0.0104). No statistically significant correlations were observed when disaggregating by intervention training group (Table 3).

**Table 3.** Correlations between changes in overall running economy (RE) and subject/training variables for all runners, Vaporfly (VP) trained runners, and traditional racing flat (FL) trained runners. Correlation p-values are corrected for multiple comparisons. Significant (<0.05) p-values are bolded

| Variables | All runners |  | VP trained runners |  | FL trained runners |  |
| --- | --- | --- | --- | --- | --- | --- |
|  | r | p | r | p | r | p |
| Change in Mass (kg) | 0.5667 | **0.0434** | 0.1867 | 0.8133 | 0.6412 | 0.0627 |
| Weekly Workout Mileage (miles) | -0.4828 | 0.5414 | 0.1465 | 0.9064 | -0.2226 | 0.5648 |
| Workout Day Soreness (1-5 rating) | 0.4012 | 0.1119 | 0.7925 | 0.4175 | -0.5400 | 0.1334 |
| Day after Workout Soreness (1-5 rating) | 0.3283 | 0.1961 | 0.7980 | 0.4118 | 0.4700 | 0.2017 |
| Weekly Mileage (miles) | -0.1961 | 0.2975 | 0.8041 | 0.4053 | 0.3180 | 0.4043 |
| Increase in Kvert | 0.1764 | 0.6259 | -0.0125 | 0.9875 | 0.1607 | 0.7610 |
| Workout Rating of Perceived Exertion (RPE) | 0.0152 | 0.9627 | -0.2943 | 0.8098 | 0.1029 | 0.7922 |

### 3.3 VP% Benefit Changes

For all subjects, when averaging across training groups, VP% benefit both in terms of RE and EC benefit, did not change from PRE (RE: 3.06 ± 1.46%; EC: 3.14 ± 1.44%) to POST (RE: 3.08 ± 1.41; EC: 3.43 ± 1.37) (RE: p = 0.4848, g = 0.0031, T = 0.0386, DF = 11; EC: p = 0.3246, g = 0.0488, T = 0.4626, DF = 9).

#### 3.3.1 FL and VP Training Group Comparison

When comparing the two training groups, however, FL and VP intervention groups displayed opposite directions of change. FL trained runners decreased VP% benefit (RE: −0.21 ± 1.03%; EC: −0.0004 ± 1.36%) whereas VP trained runners increased VP% benefit (RE: 0.53 ± 0.75%; EC: 0.97 ± 0.35%) (RE: p = 0.0872, g = 0.2319, T = 1.4521, DF = 11; EC: p = 0.0578, g = 0.2892, T = 1.7646, DF = 8) (Figure 4a and 4b). When looking at the subset of subjects who were classified as non-RFS runners during PRE (FL intervention group: n = 6; VP intervention group: n = 4), VP trained runners increased VP% benefit more than FL trained runners in terms of RE (VP: 0.53 ± 0.75%; FL: −0.22 ± 0.93%; p = 0.0762; g = 0.3060, T = 1.5377, DF = 8) and significantly more than FL trained runners when measured by EC (VP: 0.97 ± 0.35%; FL: 0.02 ± 1.15%; p = 0.0419, g = 0.4016, T = 1.9736).

**Fig. 4.**
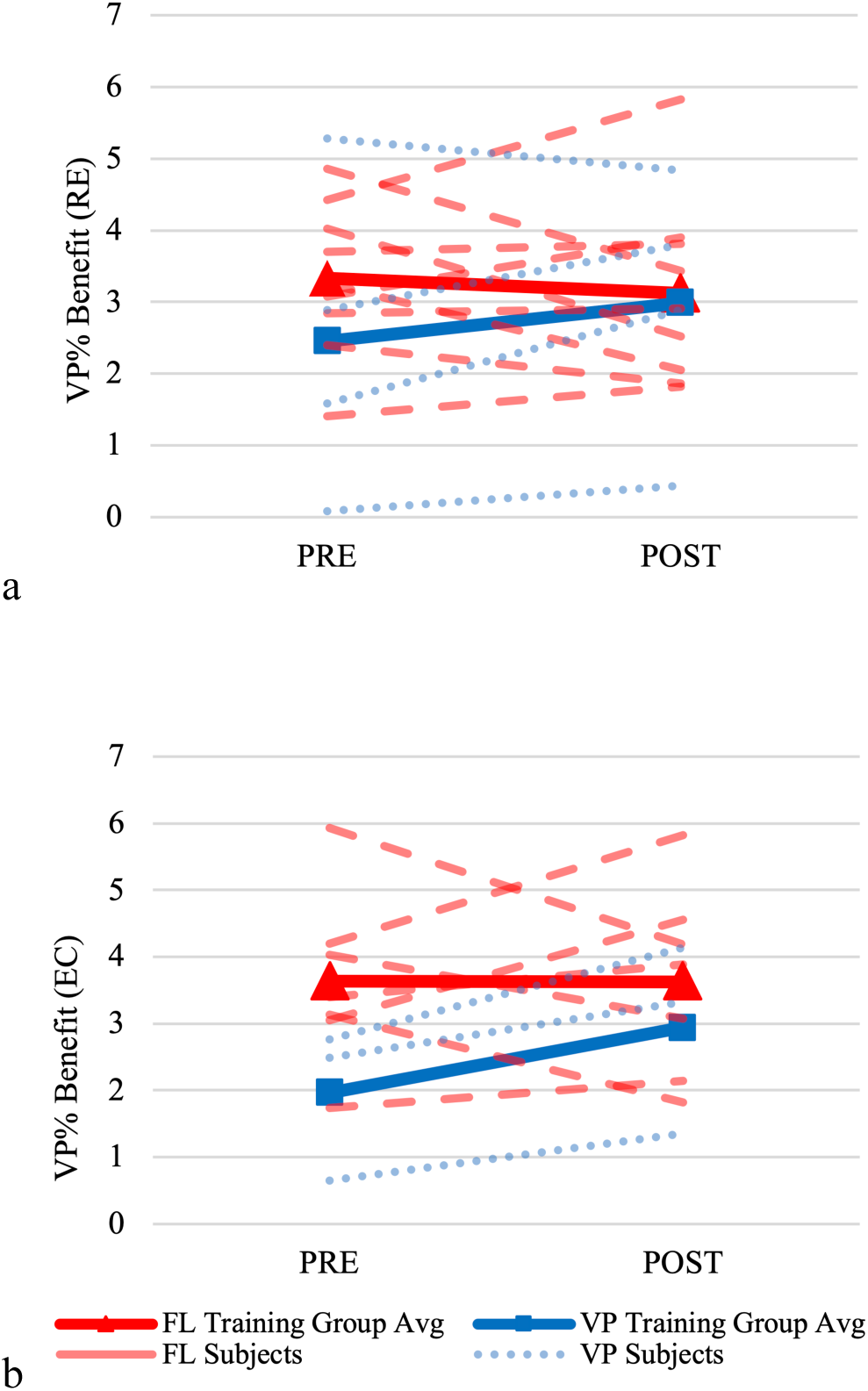
(a) Individual and group changes in acute Vaporfly performance benefits (ΔVP% benefit), in terms of (a) running economy (RE) and (b) energetic cost (EC), from pre- (PRE) to post-testing (POST)

#### 3.3.2 Associations with Changes in VP% Benefit

A total of 17 subject-based and biomechanical variables were tested in a correlational analysis with ΔVP% benefit, with variables subdivided into ankle mechanics (ΔDBS), MTP mechanics (ΔDBS), and other variables including Δsubject mass, ΔFST, ΔKvert, and steps per minute (SPM) (Table 4). When analysing all runners, ΔDBS ankle peak negative power was the only variable that was significantly associated with ΔVP% benefit (r = 0.5827; p = 0.0366) (Figure 5a). This positive association between ΔVP% benefit and ΔDBS ankle peak negative power translates to an improvement in VP% benefit as the DBS decreases. On average, FL trained runners increased peak ankle negative power from PRE to POST when running in FL and decreased from PRE to POST when running in VP, increasing the DBS in POST. VP trained runners decreased peak ankle negative power from PRE to POST in both shoes, though to a greater extent when in VP, decreasing the DBS in POST.

**Fig. 5.**
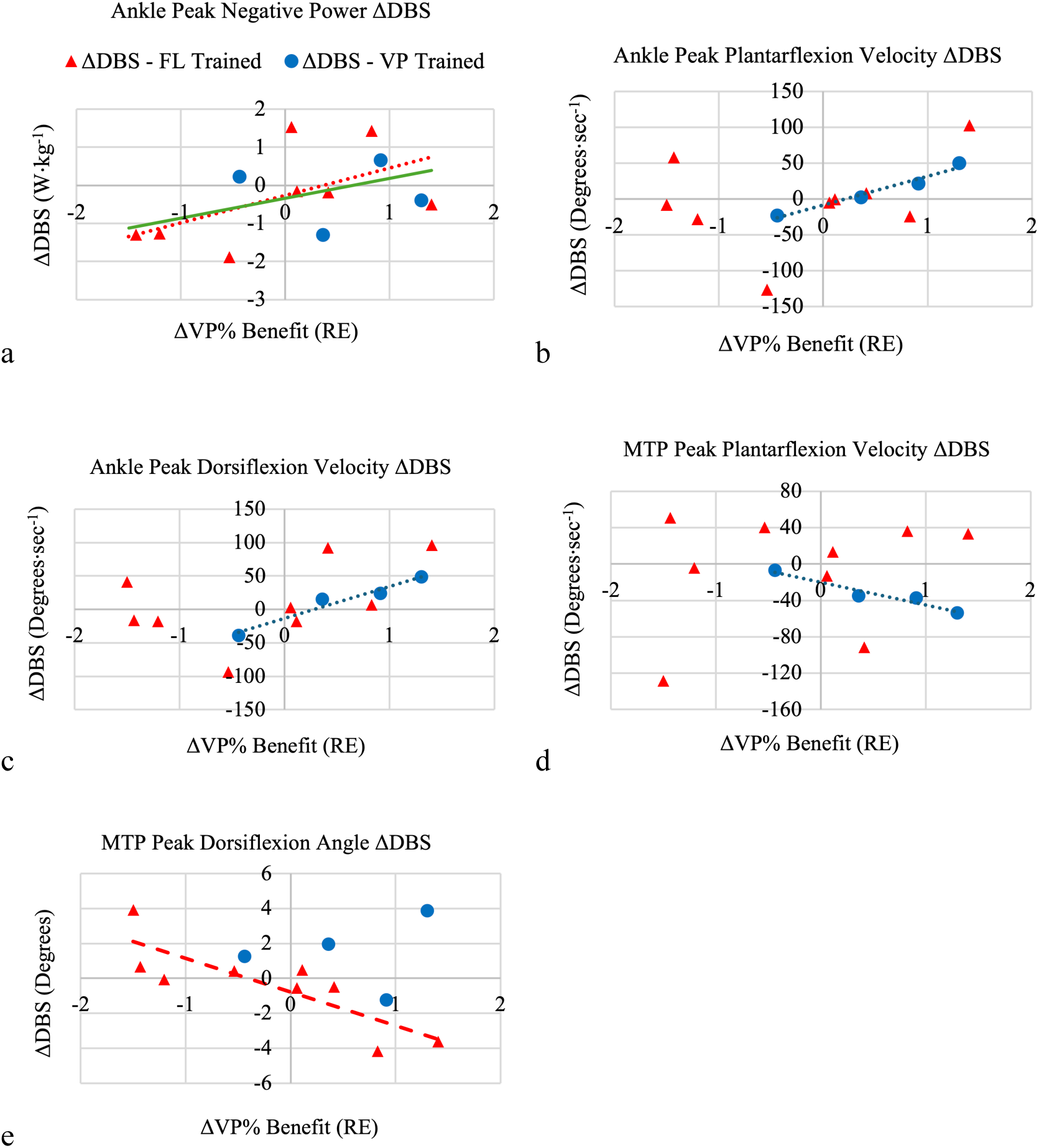
Scatterplots showing statistically significant associations between the change in acute Vaporfly (VP) performance benefits (ΔVP% benefit) and the change in the difference between shoes (ΔDBS) for biomechanical variables. Traditional racing flat (FL) and VP trained runners are represented by red triangles and blue circles respectively. Trendlines are presented for statistically significant correlations among main study runners in each graph: all runners (solid green); FL trained runners (dashed red); VP trained runners (dotted blue). Scatterplots show ΔVP% benefit and (a) ΔDBS ankle peak negative power; (b) ΔDBS ankle peak plantarflexion velocity; (c) ΔDBS ankle peak dorsiflexion velocity; (d) ΔDBS metatarsophalangeal (MTP) joint peak plantarflexion velocity; (e) ΔDBS MTP joint peak dorsiflexion angle in FL trained runners

**Table 4.** Correlations between changes in acute Vaporfly performance benefits (ΔVP% Benefit) and ankle, metatarsophalangeal (MTP) joint, and other variables of interest, among all runners, Vaporfly (VP) trained runners, and traditional racing flat (FL) trained runners. Correlation p-values are corrected for multiple comparisons. Significant (<0.05) p-values are bolded

| Variables | All runners |  | VP trained runners |  | FL trained runners |  |
| --- | --- | --- | --- | --- | --- | --- |
|  | r | p | r | p | r | p |
| Ankle Peak Plantarflexion Velocity ($\Delta DBS$) | 0.3956 | 0.1809 | 0.9835 | **0.0165** | 0.2963 | 0.4389 |
| Ankle Peak Plantarflexion Moment ($\Delta DBS$) | -0.4056 | 0.1691 | -0.8314 | 0.1686 | -0.4068 | 0.2772 |
| Ankle Peak Negative Power ($\Delta DBS$) | 0.5827 | **0.0366** | -0.0314 | 0.9686 | 0.6772 | **0.0451** |
| Ankle Peak Positive Power ($\Delta DBS$) | -0.1230 | 0.6889 | 0.0414 | 0.9586 | -0.3068 | 0.422 |
| Ankle Peak Dorsiflexion Velocity ($\Delta DBS$) | 0.5128 | 0.0731 | 0.9770 | **0.023** | 0.4700 | 0.2017 |
| Peak Foot Angle ($\Delta DBS$) | -0.0840 | 0.7850 | -0.7646 | 0.2354 | 0.0417 | 0.9153 |
| MTP Peak Dorsiflexion ($\Delta DBS$) | -0.4111 | 0.1629 | 0.1980 | 0.802 | -0.8294 | **0.0057** |
| MTP Peak Dorsiflexion Velocity ($\Delta DBS$) | -0.2377 | 0.4342 | -0.9206 | 0.0794 | -0.2047 | 0.5972 |
| MTP Peak Plantarflexion Velocity ($\Delta DBS$) | 0.0515 | 0.8674 | -0.9701 | **0.0299** | 0.2502 | 0.5161 |
| MTP Peak Plantarflexion Moment ($\Delta DBS$) | -0.3275 | 0.2747 | -0.5014 | 0.4986 | -0.3220 | 0.3981 |
| MTP Peak Positive Power ($\Delta DBS$) | 0.3893 | 0.1885 | 0.8837 | 0.1163 | 0.3875 | 0.3028 |
| MTP Peak Negative Power ($\Delta DBS$) | 0.4660 | 0.1085 | 0.3356 | 0.6644 | 0.4573 | 0.2158 |
| ΔSubject Mass | 0.4697 | 0.1054 | -0.3298 | 0.6702 | 0.6327 | 0.0674 |
| ΔFootstrike Technique | 0.3383 | 0.2583 | 0.1568 | 0.8432 | 0.6305 | 0.0687 |
| $\Delta K_{vert}$ | 0.3889 | 0.2667 | -0.6626 | 0.3374 | 0.6490 | 0.1632 |
| Steps Per Minute ($\Delta DBS$) | 0.1283 | 0.6762 | -0.7000 | 0.3 | 0.2701 | 0.4822 |

When disaggregating subjects by FL or VP intervention training group, further significant associations between ΔVP% benefit and the aforementioned variables were found (Table 4). In VP trained runners, statistically significant, strong associations were observed between ΔVP% benefit and three ΔDBS variables: (1) ankle peak plantarflexion velocity (r = 0.9835; p = 0.0165) (Figure 5b); (2) ankle peak dorsiflexion velocity (r = 0.9770; p = 0.0230) (Figure 5c); and (3) MTP joint peak plantarflexion velocity (r = −0.9701; p = 0.0299) (Figure 5d). In FL trained runners, two statistically significant correlations were observed between ΔVP% benefit and two ΔDBS variables: (1) ankle peak negative power (r = 0.6772; p = 0.0451) (Figure 5a); and (2) MTP joint peak dorsiflexion angle (r = −0.8294; p = 0.0057) (Figure 5e). See Online Resource 1 for correlational matrices between ΔVP% benefit and biomechanical variables of interest among all runners, VP trained runners, and FL trained runners.

Here, the directions of these associations are explained. The positive association between ΔVP% benefit and peak ankle negative power (ΔDBS) was similar in FL trained runners as seen among the full sample of runners. In this association, runners who increased the DBS in peak ankle negative power over the course of the intervention decreased their VP% benefit more. In VP trained runners, instead of ankle power, peak ankle plantarflexion and dorsiflexion velocities (ΔDBS) were strongly correlated with ΔVP% benefit. VP trained runners increased the DBS in peak ankle plantarflexion velocity by increasing peak ankle plantarflexion velocity when running in FL and decreasing velocity when running in VP (Table 5), and the greater this DBS, the greater the observed VP% benefit. Also, in VP trained runners, the DBS in peak ankle dorsiflexion velocity decreased, with runners having a faster peak ankle dorsiflexion velocity when running in VP and slower in FL from PRE to POST. Greater VP% benefit was associated with a smaller DBS in peak ankle dorsiflexion velocity in POST for VP trained runners.

**Table 5.** Biomechanical variables for all runners, divided by training intervention group, in pre- (PRE) to post-testing (POST). Values are shown for Vaporfly (VP) and traditional racing flat (FL) trials in PRE and POST

| | FL Training Group (Mean ± SD) | | | | VP Training Group (Mean ± SD) | | | |
| --- | --- | --- | --- | --- | --- | --- | --- | --- |
|  | PRE |  | POST |  | PRE |  | POST |  |
|  | FL | VP | FL | VP | FL | VP | FL | VP |
| Peak Ankle Plantarflexion Velocity (degrees·sec<sup>-1</sup>) | 889.3 ± 106.2 | 901.5 ± 85.8 | 868.6 ± 95.9 | 883.5 ± 111.1 | 914.9 ± 136.7 | 907.8 ± 65.4 | 920.7 ± 123.9 | 901.0 ± 70.3 |
| Peak Ankle Dorsiflexion Velocity (degrees·sec<sup>-1</sup>) | -555.3 ± 148.0 | -481.3 ± 164.3 | -514.7 ± 114.0 | -451.2 ± 132.6 | -527.2 ± 134.7 | -464.3 ± 103.8 | -523.1 ± 154 | -472.5 ± 128.5 |
| Peak MTP Dorsiflexion Angle (degrees) | -22 ± 3.2 | -10.9 ± 3.9 | -23 ± 2.4 | -11.5 ± 3.0 | -21.9 ± 2.9 | -10.5 ± 3.9 | -21.9 ± 2.5 | -12.1 ± 2.9 |
| Peak MTP Dorsiflexion Velocity (degrees·sec<sup>-1</sup>) | -505.4 ± 64.5 | -256.4 ± 50.7 | -504.5 ± 55.3 | -266.7 ± 52.0 | -492.2 ± 63.2 | -283.4 ± 59.6 | -488.9 ± 65.4 | -285 ± 63.7 |
| Peak MTP Plantarflexion Velocity (degrees·sec<sup>-1</sup>) | 683.1 ± 103.6 | 371.2 ± 88.3 | 682.9 ± 64.3 | 378.0 ± 86.7 | 773.5 ± 135.7 | 418.9 ± 112.4 | 744.8 ± 146.1 | 423.4 ± 112.5 |
| Peak Ankle Plantarflexion Moment (Nm·sec<sup>-1</sup>) | 5.0 ± 0.9 | 4.9 ± 0.8 | 4.9 ± 0.9 | 4.8 ± 0.9 | 5.2 ± 0.1 | 5.1 ± 0.2 | 5.2 ± 0.2 | 5.1 ± 0.2 |
| Peak MTP Plantarflexion Moment (Nm·sec<sup>-1</sup>) | 2.4 ± 0.4 | 2.5 ± 0.4 | 2.4 ± 0.5 | 2.5 ± 0.5 | 2.6 ± 0.2 | 2.7 ± 0.2 | 2.6 ± 0.2 | 2.7 ± 0.3 |
| Peak Ankle Negative Power (W·sec<sup>-1</sup>) | -10.9 ± 3.2 | -8.8 ± 3.2 | -11.0 ± 3.1 | -8.6 ± 3.1 | -11.0 ± 1.5 | -9.2 ± 0.4 | -10.9 ± 1.3 | -8.9 ± 0.9 |
| Peak Ankle Positive Power (W·sec<sup>-1</sup>) | 54.0 ± 16.6 | 45.4 ± 13.1 | 50.7 ± 15.8 | 43.3 ± 12.1 | 55.3 ± 10.9 | 47.0 ± 5.2 | 56.3 ± 10.6 | 45.4 ± 7.5 |
| Peak MTP Positive Power (W·sec<sup>-1</sup>) | 9.4 ± 5.2 | 7.1 ± 3.2 | 7.9 ± 4.4 | 6.5 ± 2.5 | 11.9 ± 5.1 | 8.3 ± 1.9 | 10.2 ± 5.1 | 7.7 ± 2.9 |
| Peak MTP Negative Power (W·sec<sup>-1</sup>) | -19.1 ± 5.3 | -10.6 ± 3.6 | -19.3 ± 5.3 | -11.2 ± 3.7 | -19.6 ± 2.1 | -11.8 ± 2.1 | -19.1 ± 3.3 | -11.8 ± 2.7 |
| Average Steps Per Minute | 182.6 ± 12.6 | 180.9 ± 12.7 | 183.2 ± 13.7 | 182.8 ± 14.0 | 183.9 ± 6.5 | 182.7 ± 7.6 | 183.6 ± 7.4 | 182.4 ± 6.8 |
| Foot Angle at Contact (degrees) | 1.9 ± 3.3 | 1.9 ± 3.7 | 1.5 ± 3.1 | 1.7 ± 3.0 | 5.4 ± 2.5 | 7.3 ± 4.2 | 2.0 ± 2.0 | 1.3 ± 1.9 |

While no MTP joint ΔDBS variables were associated with ΔVP% benefit among all runners, in VP trained runners only, ΔVP% benefit was negatively associated with ΔDBS MTP joint peak plantarflexion velocity. VP trained runners, on average, decreased MTP joint peak plantarflexion velocity when running in FL and increased velocity when running in VP, leading to a decreased DBS in POST, and in this case the lesser the DBS, the greater the observed VP% benefit. Overall, VP trained runners had a greater decrease in the MTP joint peak plantarflexion velocity DBS compared to FL trained runners (Table 5).

Finally, VP trained runners decreased MTP joint peak dorsiflexion velocity when running in FL and increased velocity when running in VP, also decreasing the DBS in POST but not to as great of an extent as FL trained runners. The strong negative correlation between ΔVP% benefit and ΔDBS MTP joint peak dorsiflexion velocity observed in VP trained runners means that a decrease in the DBS in POST is associated with greater VP% benefit. In FL trained runners, ΔDBS peak MTP dorsiflexion angle was strongly negatively correlated with ΔVP% benefit, meaning a greater DBS is associated with a higher VP% benefit. On average, FL trained runners increased the DBS in peak MTP joint dorsiflexion from PRE to POST (Table 5).

## 4 Discussion

### 4.1 Intervention Characteristics

Of 24 runners that started the training intervention, 13 completed the training intervention, which is a low retention rate. Of the 11 runners who did not complete the study, 7 runners withdrew for reasons not related to injury (e.g., college stress, scheduling conflicts, sickness, reluctance to use FL in workouts, and discontinued communication). Two subjects withdrew due to injuries unlikely related to intervention footwear use: one injury occurred before the participant’s first intervention workout and the other runner withdrew due to a mild ankle sprain prior to POST.

Adjustments to intervention footwear used and break-in periods markedly improved the attrition rate in FL assigned runners compared to the pilot study [25]. In the present study, four runners assigned to the FL intervention group withdrew from the study – two due to complaints related to their use of FL and two did not cite adverse effects of FL for their reasons for withdrawal. Of the two who withdrew for footwear-related reasons, one runner withdrew after the first workout, citing aggravation of their calf and not breaking in FL as suggested. The second runner self-reported Achilles tightness that accumulated between successive workouts and races in FL. These calf and Achilles complaints for FL trained runners are in line with the injuries commonly reported in other studies investigating the effects/risks of transitioning to barefoot or minimalist footwear, who have found increased risks of plantarflexor injuries, among other injuries, such as metatarsal stress fractures [16, 49]. Compared to the pilot study, where 3 of 5 FL assigned runners discontinued training in FL due to complaints of muscular soreness [25], the present study addressed this issue by selecting a more protective traditional racing flat (Nike Zoom Rival Waffle 5), compared to the Nike Victory Waffle 5, for FL intervention group runners and encouraging a break-in period.

Attrition rates from Matties et al. [25], coupled with a lack of footwear related injuries, greater intervention footwear adherence rates, lower average perceived exertion, and lower average muscular soreness reported in VP trained runners in both the pilot and present studies, suggest that runners prefer training in VP. The potential recovery advantages afforded by VP in interval workouts [17, 50], along with other hard workouts completed by our runners, seems to be a common factor driving this preference. This is further supported by the self-selection of FL assigned runners into the VP training group in the pilot study [25].

### 4.2 Overall ME Changes and Function Tests

While the specific effects of a season of cross-country training, including endurance and strength training, on ME are relatively unknown [51], we did see a small improvement in overall ME on average for participants that completed the intervention. This agrees with previous literature discussing the influence of endurance training on ME [51–54]. On average, a non-significant increase in Kvert was observed, which may partially explain an increase in overall ME with previous research showing an increase in leg/vertical stiffness coinciding with an improved ME [55–59], though findings are mixed [60].

Comparing overall ME between VP and FL trained runners, we see that FL trained runners tend to improve overall ME slightly more than VP trained runners, with no statistically significant difference between training groups. The same pattern is observed when combining RE and EC data from the pilot study and the second, present study phase. It should be noted that the between-day reliability of ME measurements are affected by a multitude of extraneous factors, such as time of day, diet, and fatigue, therefore these results should be interpreted with caution [61]. These changes were similar to changes in RE measured by minimalist footwear interventions, who saw non-significantly greater RE improvements in runners who trained in the minimalist footwear condition [19–21]. Though, with increased soreness, RPE, and self-reported injuries, the pilot and present studies show that the potential higher risks associated with training in FL may limit these marginal performance gains to those who are able to tolerate training in a more minimalist shoe [25].

### 4.3 Explaining Overall ME Changes

Among all runners in the present study, an increase in runner mass was associated with better RE, likely driven by the unitary definition of RE: ml·kg^-1^·min^-1^. In VP trained runners, stronger positive associations were observed between overall RE improvement and greater weekly workout mileage, workout day soreness, and day after workout soreness (Table 3). Although these associations were non-significant, they were either stronger, or, for weekly workout mileage, in an opposite direction than in FL trained runners. The fact that mileage run, and soreness attained, were associated with RE improvement in VP trained runners, but not FL trained runners or all runners (in fact, mileage was negatively associated with change in RE in FL trained runners), suggests that training in VP allows runners to better take advantage of workout mileage and soreness to achieve gains in RE. These findings should be interpreted with caution due to the small sample size of the present study.

### 4.4 VP% Benefit Changes

Acute RE and EC responses to VP compared to FL (VP% benefit), were about 3% on average among subjects in this sample, which is in line with the existing literature showing acute responses averaging between 2.5 to 4% [6–8, 37]. While closely missing statistical significance in this small sample, on average FL trained runners decreased VP% benefit slightly and VP trained runners increased VP% benefit, with a greater magnitude of VP% benefit change for VP trained runners. This finding supports the footwear specificity of training principle hypothesized by Matties et al. [25]. When considered together, these two studies have oppositely balanced FL and VP training group sizes and provide similar evidence for this footwear specificity of training principle.

Based on that pattern observed in the pilot study where non-RFS VP trained runners improved VP% benefit more than RFS runners [25], it would be hypothesized that all VP-trained runners in the present study, who all happened to be non-RFS runners, would increase their VP% benefit. Supporting this hypothesis, in the present study 3 of 4 VP trained runners increased VP% benefit in terms of RE and all runners included in EC analysis (3 of 3) increased VP% benefit. Evidence from both studies suggests that training in VP increases VP% benefit, contrary to training in FL, with non-RFS runners potentially benefiting to a greater extent by transitioning towards a RFS pattern when running in VP.

### 4.5 Explaining VP% Benefit Changes

ME data show that training in VP seems to increase runners’ acute VP% benefit. The correlational analysis of runners in the VP and FL training groups reveals a significant association between decreasing the ankle peak negative power DBS and thereby increasing VP% benefit. Contrary to this finding, previous research comparing the biomechanics of runners wearing AFT, shows decreased peak ankle positive and negative power in the AFT condition, suggesting a lower storage of energy in the triceps surae [12]. This would suggest that the DBS should be greater to observe a greater VP% benefit.

When further breaking down the correlation analysis to investigate relationships within each training group, we see that FL trained runners follow the trend we see in all runners (FL and VP trained runners), with a decreased ankle peak negative power DBS being associated with an increased VP% benefit, but there is no association between these variables in VP trained runners. This suggests that changing peak ankle negative power is likely not a mechanical adaptation that VP trained runners are altering to improve VP% benefit, though may be a characteristic that remains present when acutely comparing shoes.

Instead, in VP trained runners, increased ankle plantarflexion velocity DBS and decreased ankle dorsiflexion velocity DBS were associated with increased VP% benefit. On average VP trained runners increased ankle plantarflexion velocity DBS with higher peak ankle plantarflexion velocity when running in FL and slower when running in VP. One reported mechanism through which AFT potentially improves running economy acutely is through slower ankle plantarflexion velocities, which allow for more metabolically efficient force production [13]. Given the tendency of VP trained runners to increase ankle plantarflexion velocity DBS, particularly by further slowing plantarflexion velocity when in VP, the acute metabolic and mechanical differences reported by others comparing AFT and FL support this finding [7, 12, 13]. This is one potential mechanism of habituation with VP use that allows for increased VP% benefit in VP trained runners.

A significant association was also observed in VP trained runners between decreased peak ankle dorsiflexion velocity DBS and increased VP% benefit. VP trained runners decreased peak ankle dorsiflexion velocity DBS in POST by decreasing peak dorsiflexion velocity running in FL and increasing when running in VP. This decreased DBS in POST for VP trained runners further supports the increased VP% benefit observed based on the correlational analysis. A similar decrease in peak ankle dorsiflexion velocity DBS was observed in FL trained runners but was achieved by slower velocities when running in VP and FL in POST, with a greater decrease in dorsiflexion velocity observed in FL. Interestingly, this change in the DBS for FL trained runners was not associated with VP% benefit. While previous studies have shown that AFT acutely does not seem to significantly impact peak ankle dorsiflexion velocity [12], subjects in the present study, on average, had a slower peak ankle dorsiflexion velocity in VP during both lab sessions.

One possible interpretation of these results could be that VP trained runners are increasing ankle dorsiflexion velocity when running in VP to transition anteriorly through the shoe more quickly in the first 15-25% of stance. Matijevich et al. [62] suggests that more optimal timing of AFT energy storage and return may allow for runners to further increase metabolic savings. With VP trained runners in the present study increasing ankle dorsiflexion velocity, these runners may be taking better advantage of energy returned in the heel region of the shoe, which is known to be returned at a much higher frequency than optimal in stance phase due to the characteristics of polyether block amide (PEBA) midsole foam [11]. Considering that two of the VP trained runners transitioned to a RFS in POST, this would allow them to potentially interact with more of the foam in the heel of VP. Given the lack of previous literature establishing a relationship between AFT and peak ankle dorsiflexion velocity, it is difficult to determine potential mechanisms through which changes in this measure may impact changes in VP% benefit.

Further distally, a significant association was observed in VP trained runners between decreased MTP joint peak plantarflexion velocity DBS and increased VP% benefit. Both VP and FL trained runners decreased MTP joint peak plantarflexion velocity DBS in POST, though the VP trained runners did so to a greater extent. With MTP joint peak plantarflexion velocity being slower in VP compared to FL acutely, VP trained runners decreased the DBS by consistently decreasing peak MTP joint plantarflexion velocity when running in FL (Figure 5d), whereas FL trained runners maintained the same velocity when running in FL and increased when running in VP. Acute decreases in MTP joint plantarflexion velocity are commonly shown in research looking at AFT or footwear with a stiffening plate, with the stiffening agent acting in parallel with the intrinsic foot muscles to help aid in stiffening the MTP joint during push off, decreasing energy lost at the MTP joint [13]. In VP trained runners, the decrease in MTP joint plantarflexion velocity when running in FL was greater than the increase in VP during POST. It is difficult to identify a specific mechanism through which training in VP may cause this specific mechanical adaptation and influence VP% benefit, though this may suggest that VP trained runners may become more reliant on the embedded carbon plate to help stiffen their foot after training in VP. When these runners were tested in FL during POST, a decreased MTP joint plantarflexion velocity was observed in FL, potentially due to a lower contribution of the intrinsic foot muscles. While the mechanisms of this change remain unclear, particularly with how a decrease in MTP peak plantarflexion velocity DBS is associated with an increased VP% benefit, it seems that VP trained runners may increasingly rely on the stiffening plate during toe off to stiffen their MTP joint passively, further reducing the metabolic demand on the intrinsic foot muscles, which may serve to increase VP% benefit.

In FL trained runners, a significant association was observed between increased MTP joint peak dorsiflexion angle DBS and increased VP% benefit. On average, FL trained runners increased MTP joint peak dorsiflexion angle DBS in POST, mainly by increasing MTP joint peak dorsiflexion to a greater extent when running in FL. On the other hand, VP trained runners increased peak MTP joint dorsiflexion to a greater extent when running in VP, and this measure was not significantly correlated with changes in VP% benefit. In agreement with previous studies, runners in the present study exhibited smaller peak MTP dorsiflexion angles when running in VP compared to FL. Thus, the acute increase in MTP joint dorsiflexion seen in FL relative to VP seems to translate to greater peak MTP joint dorsiflexion angles, particularly when running in FL, for FL trained runners after using FL in workouts. There are multiple potential explanations for how this may translate to increased VP% benefit in FL trained runners. It is possible that the relatively greater MTP joint dorsiflexion in FL during POST translates to more energy lost at the MTP joint, which does occur but to a greater degree when FL trained runners run in VP during POST. Alternatively, the potentially greater range of motion at the MTP joint could increase metabolic demand when running in FL by requiring force production by the intrinsic foot muscles over a longer duration or larger range of motion, improving VP% benefit by increasing the relative energetic cost in FL.

### 4.6 Practical Considerations for Workout Footwear Choice

One of the main motivations of the current research was to inform runners, coaches, and footwear companies about the potential benefits and drawbacks of training in AFT. The present study focused on collegiate and elite club cross-country runners, who train on a variety of surfaces that may not be common for other runners, but the results of this study still provide insightful data to support beliefs likely already held by runners and coaches and may suggest ways to optimize workout footwear choice. A main finding of the pilot study was the specificity of training principle with running workout footwear, with runners improving their relative RE to a greater extent in the shoe they trained in for workouts over the 8-week intervention [25]. The present study further supports this hypothesis, with the magnitude of VP% benefit increase for VP trained runners being greater than the VP% benefit decrease seen in FL trained runners. This suggests that this potential footwear specificity of training principle may have more potential for VP trained runners to habituate to VP effectively, increasing VP% benefit by training in the shoe prior to race day.

It should be considered that increasing VP% benefit by training in VP may not be advantageous for all runners. Some may be training for a race that AFT is not an option for or they prefer a different shoe type. For these runners, results from the present study may provide a rationale to use their planned racing footwear during running workouts. At a more fundamental level, this racing footwear habituation may allow for increased comfort, which has been shown to potentially improve RE when comparing different footwear of varying comfort levels, though findings are mixed [63, 64].

Changes in shoe-specific efficiency may help contribute towards better performance on race day, though overall changes in ME due to training likely have a larger role for the majority of runners. The pilot and present studies show that FL trained runners may have slightly higher improvements in overall ME, though these runners consistently report higher RPE, muscular soreness, and more frequently self-report musculoskeletal pain. This trade off should be considered, along with the potential for VP to increase runners’ ability to tolerate a higher training workload. Training in VP may allow runners to increase their workout volume, intensity of workouts, and/or workout frequency. Runners and coaches likely already intuitively know there is potential for higher workload when training in AFT, exemplified by the increased popularity of workouts that take advantage of these benefits, such as double threshold workouts.

The present study design does not allow for determination of the minimum period of habituation to VP or FL to elicit improvements in shoe-specific economy. For those who would like to avoid using more expensive, less durable footwear regularly in training, many footwear companies that produce AFT also create a “training companion” that is more durable and employs similar technologies (e.g., Nike Zoom Fly 5, Saucony Endorphin Speed 4, Adidas Boston 12). These models may allow for similar training adaptations and responses, as training in AFT would, while improving workout footwear longevity and decreasing cost. Though the effects of these types of workout shoes are unknown at this time, if the footwear specificity of training principle applies, then these shoes may provide a great opportunity to closely mimic running in AFT during workouts and provide a degree of habituation.

### 4.7 Limitations

The results of the present study should be interpreted conservatively, however, due to a variety of limitations. Runners were included from a variety of collegiate and club cross-country programs, meaning training was less standardized than the pilot study, where runners belonged to the same collegiate program [25]. This should also be considered when it comes to strength training, which is suggested to positively influence RE [65]. Differences in training protocols between participants may have influenced measures of overall ME. The measure of overall ME should be interpreted cautiously, particularly with a smaller sample size, due to the impact of many different factors on this measure, including participant fatigue, diet, time of day, among other factors [61].

Associations were tested between metabolic and biomechanical data, though these data were collected at separate running speeds. With participants’ training being targeted for cross-country races during the intervention period, running mechanics were collected at their cross-country race pace to more accurately capture potential habituations effects. RE and EC measures require submaximal testing speeds [61], limiting VP% benefit measurement to the 5-minute submaximal trials. A limitation of the present methodology and associations tested are that runners may change footwear-specific mechanics at faster paces in different ways than at slower paces.

The intervention attrition rate was improved in the present study compared to the pilot study, though attrition was still high due to a variety of participant-specific factors. This high attrition resulted in a small sample size that was unbalanced, with relatively more FL assigned runners completing the intervention. While the present study does support the main results from Matties et al. [25], there are some differences between shoe models used, data collection protocols, metabolic carts used, and most importantly a lack of true random assignment in the pilot study due to self-selection. However, the oppositely balanced intervention group sizes and similarities between footwear conditions in the current study compared to Matties et al. [25], along with the same findings that support a footwear specificity of training principle, provide evidence that habituation may influence footwear-specific performance.

Within intervention groups, runners were requested to maintain at least an 80% adherence rate, or complete 80% of their hard running workouts, in their assigned intervention shoe. This was done to increase athlete and coach willingness to participate in the study during a competitive season, allow for a break-in period, and reduce attrition rates. Over the 8-week period, the 80% adherence rate allowed for three hard running workouts in an alternate shoe. FL trained runners avoided AFT during the intervention. Even with about a 98% adherence rate in the VP intervention group, it is recognized that only using intervention footwear twice a week is not as frequent as other footwear intervention studies, though the total mileage accumulated in intervention footwear was similar or greater than other footwear interventions [19–21, 66]. In this sample of collegiate and competitive club cross-country runners, this twice per week frequency and weekly mileage completed in the intervention footwear matched real workout footwear usage patterns based on runner’s intake questionnaire responses.

Biomechanical data were limited to the sagittal plane and may have been affected by small differences in marker placement between testing sessions. Furthermore, the present analysis is limited to group averages, though individual analyses of runners’ mechanics and shoe-specific responses may provide further clarity on adaptations being made to increase VP% benefit or overall ME.

The present study was also limited to about an 8-week intervention length based on constraints surrounding the collegiate cross-country season and athlete availability for lab testing. While the length of this intervention was long enough to create adaptations based on previous footwear intervention literature [14, 22], a longer intervention period may allow for more significant differences to be observed.

### 4.8 Recommendations for Future Research

Additional research should be conducted to further understand how long-term AFT use influences training adaptations, recovery, injury, and race day performance. Future analyses may be conducted to further explore how changes in lower limb coordination patterns, electromyography activation, and other gait characteristics, impact overall ME and VP% benefit. Populations of other runners should be considered, as well as different footwear models and technologies. With workout footwear designs changing dramatically in the last few years, more of these shoes incorporate AFT characteristics. Future research using different control shoes, such as a non-plated workout shoe that still uses supercritical foam technology or a plated EVA foam shoe, may provide data that is more applicable to the majority of runners given the characteristics of today’s workout footwear choices.

The potential footwear specificity of training principle suggested by the present and Matties et al. [25] should be further investigated. Schwalm et al. [24] observed no habituation effect on RE or EC with participants between different AFT models and their own personal pair(s) of AFT with at least 20 km of use. It should be considered that this comparison was made between habituated and non-habituated AFT models, likely producing smaller shoe-specific ME differences than the VP and FL models used in the present study. Similar interventions conducted with AFT, track spikes, and trail running footwear, may support or refute this footwear habituation hypothesis. Alternatively, a larger sample of runners and more advanced statistical analyses with a similar intervention protocol as the present study may provide evidence to explain why runners experience varying acute metabolic responses to AFT. This may allow footwear companies to better understand individual runners’ characteristics and their impact on acute and long-term AFT responses, allowing for further individualization of footwear technologies to maximize performance benefits for runners.

## 5 Conclusions

This research investigated the effects of training in VP compared to FL on overall metabolic efficiency (ME), acute VP performance benefits (VP% benefit), and mechanical adaptations that are associated with changes in shoe-specific efficiencies. A footwear specificity of training principle was observed, with VP trained runners improving their VP% benefit and FL trained runners decreasing their VP% benefit in POST. The biomechanical analyses suggest that VP trained runners experience footwear-specific biomechanical adaptations that further increase their VP% benefit. These mechanical adaptations include decreased ankle plantarflexion velocity in VP, increased ankle dorsiflexion velocity in VP, and decreased MTP joint plantarflexion velocity in FL.

No significant differences in overall ME improvement from PRE to POST were observed between FL and VP training groups, with a considerable amount of inter-individual variability highlighting limitations in this measure. With small overall ME differences between groups observed, training in FL may afford potentially greater overall ME improvements from workouts for some individuals, but the potentially higher risk of injury and lower training tolerance should be considered. The present study and the pilot study presented by Matties et al. [25] show that, on average, VP trained runners reported lower, though not significantly different, ratings of perceived exertion (RPE) and experienced less perceived muscular soreness both the day of and day after a workout/race. This suggests that an advantage of training in VP is increased runner capacity for workout volume, intensity, or frequency.

Future research investigating the minimum exposure for biomechanical adaptations, or habituation, to occur in response to training footwear would inform runners and coaches about when to introduce race day footwear into training cycles. Identifying potential mechanisms of habituation to AFT, particularly on an individual level, may also provide insight for researchers and the footwear industry into the mechanical characteristics that can be further influenced to improve acute AFT performance benefits.

## Declarations

### Funding

This work was supported by the Nike Sport Research Lab through a footwear grant associated with the 2023 Nike Award for Athletic Footwear Research: Celebrating inclusivity in footwear science; the California State University East Bay College of Education and Allies Studies; the California State University East Bay Center for Student Research, and the California State University East Bay Kinesiology Research Group.

### Conflicts of Interest

Footwear for this research was partially provided by Nike based on proposal selection for the 2023 Nike Award for Athletic Footwear Research. Nike did not have a role in the execution of this research, including data collection, analysis, or interpretation of results.

### Data Availability

The anonymized dataset for this study is available in Online Resource 2.

### Ethics Approval

This project was approved by the California State University, East Bay Institutional Review Board (CSUEB-IRB-2022-150). The study was performed in accordance with the standards of ethics outlined in the Declaration of Helsinki.

### Consent to Participate

All participants provided written consent.

### Consent for Publication

Not applicable.

### Code Availability

Not applicable.

### Author Contributions

J.M. and M.R. conceptualized this research project, designed methodology, acquired funding, and collected data. All authors conducted formal analyses and interpreted the results. J.M. and M.R. drafted the original manuscript. All authors reviewed and edited the manuscript prior to submission.

## Supporting information

Online Resource 1 - Supplemental Figures

Online Resource 2 - Supplemental Dataset

## Acknowledgments

The authors would like to thank Vanessa Yingling, PhD, FACSM, and Albert Mendoza, PhD, for their feedback and guidance over the course of this research. The authors would also like to thank the undergraduate and graduate students who assisted with data collections and processing, including Dylan Lago, Rachael Decker, Karamvir Singh, and Ronaldo Flores, as well as the East Bay lab coordinator, Emily Van Horn. Finally, the authors would like to thank Dustin Joubert, PhD, and Kim Hébert-Losier, PhD, for their contributions and fruitful discussions, Coach Jordan Rodriguez for supporting athlete participation in our pilot phase, and the athletes who participated in this research.

