## Supplementary material for "Footwear-specific biomechanical and energetic responses to 8 weeks of training in advanced footwear technology": Online Resource 1 - Supplemental Figures

Journal Preprint

**Supplemental Figures**

Figure S1. Correlation matrix showing the magnitude and direction of associations between changes in acute Vaporfly running economy improvements (ΔVP% benefit) and changes in biomechanical differences between shoes (ΔDBS), at the ankle, among all runners from pre- to post-intervention. The magnitude and direction of each Pearson correlation (r) is illustrated by the colour and size of each circle, detailed by the vertical axis on the right. Non-significant p values are noted within each circle. Significant associations ($p < 0.05$), noted with a bracketed number, were: [1] p = 0.0366; [2] p = 0.0034; [3] p = 0.0456; [4] p = 0.0253.

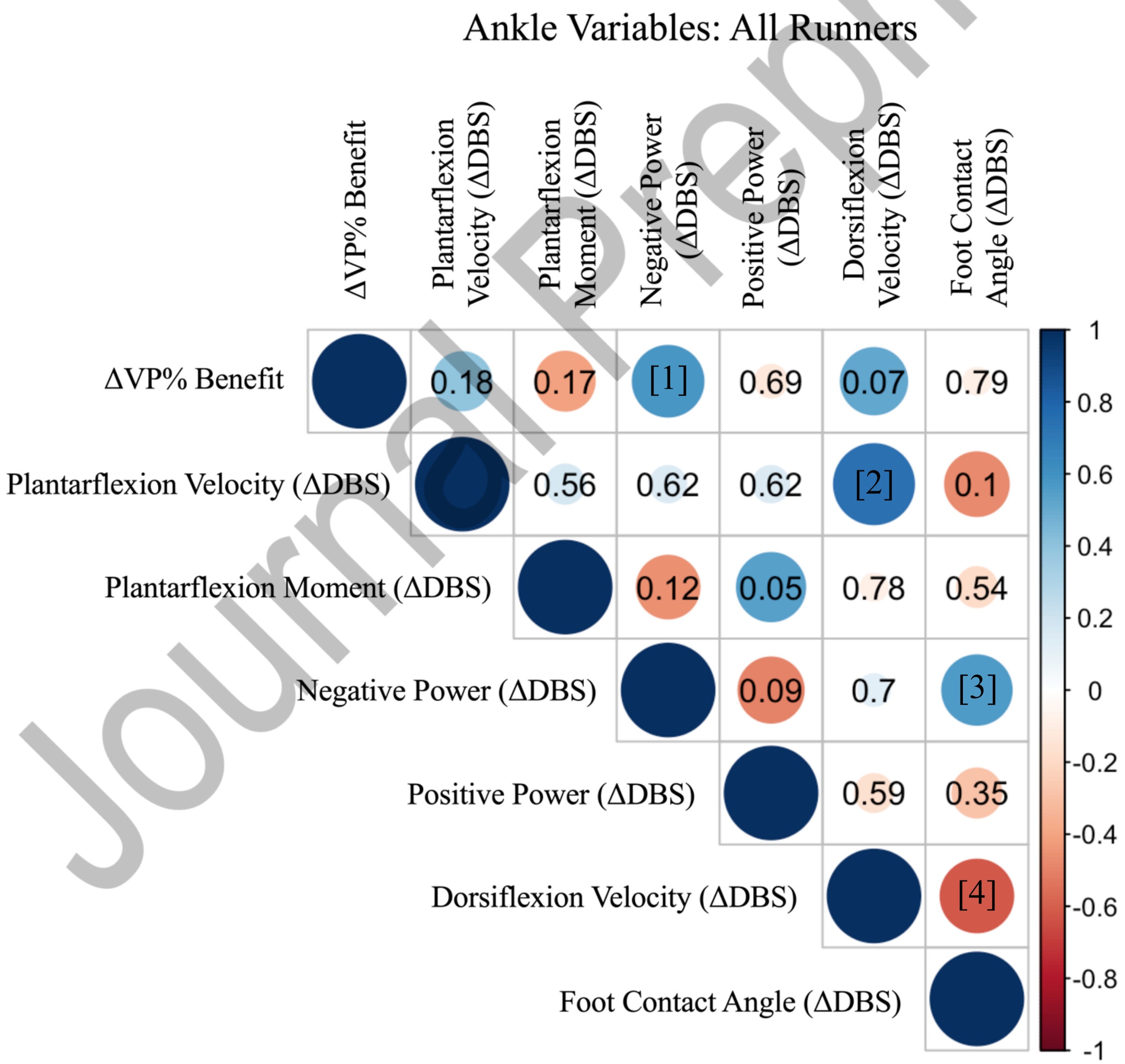


Journal Pre-proof

Figure S2. Correlation matrix showing the magnitude and direction of associations between changes in acute Vaporfly running economy improvements (ΔVP% benefit) and changes in biomechanical differences between shoes (ΔDBS), at the metatarsophalangeal (MTP) joint, among all runners from pre- to post-intervention. The magnitude and direction of each Pearson correlation (r) is illustrated by the colour and size of each circle, detailed by the vertical axis on the right. Non-significant p values are noted within each circle. Significant associations ($p < 0.05$), noted with a bracketed number, were: [1] $p = 0.0454$; [2] $p = 0.0170$.

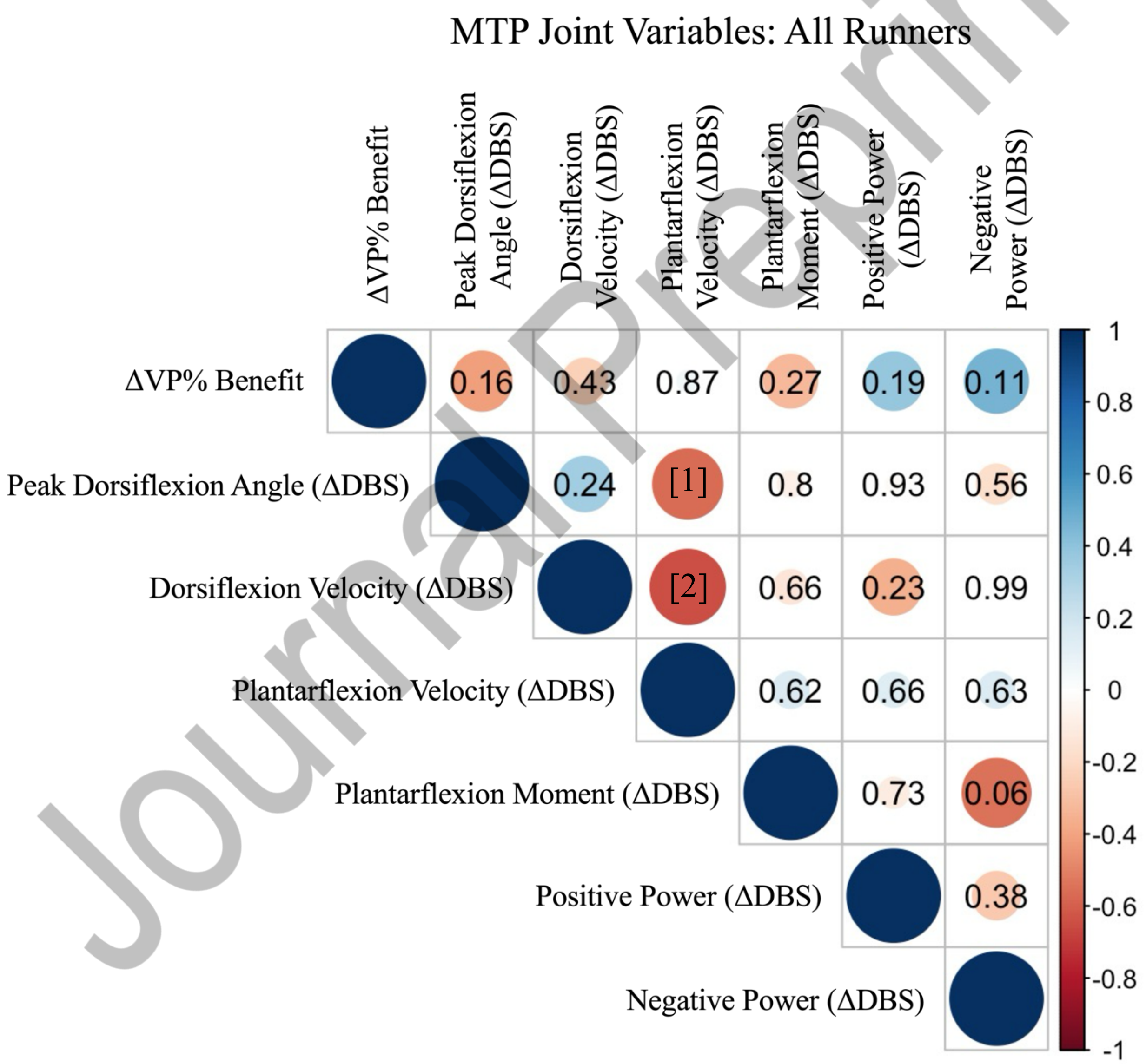


Journal Pre-proof

Figure S3. Correlation matrix showing the magnitude and direction of associations between changes in acute Vaporfly running economy improvements (ΔVP% benefit) and additional variables of interest, among all runners from pre- to post-intervention. The magnitude and direction of each Pearson correlation (r) is illustrated by the colour and size of each circle, detailed by the vertical axis on the right. Non-significant p values are noted within each circle. Significant associations ($p < 0.05$), noted with a bracketed number, were: [1] p = 0.0006; [2] p = 0.0001; [3] p = 0.0323.

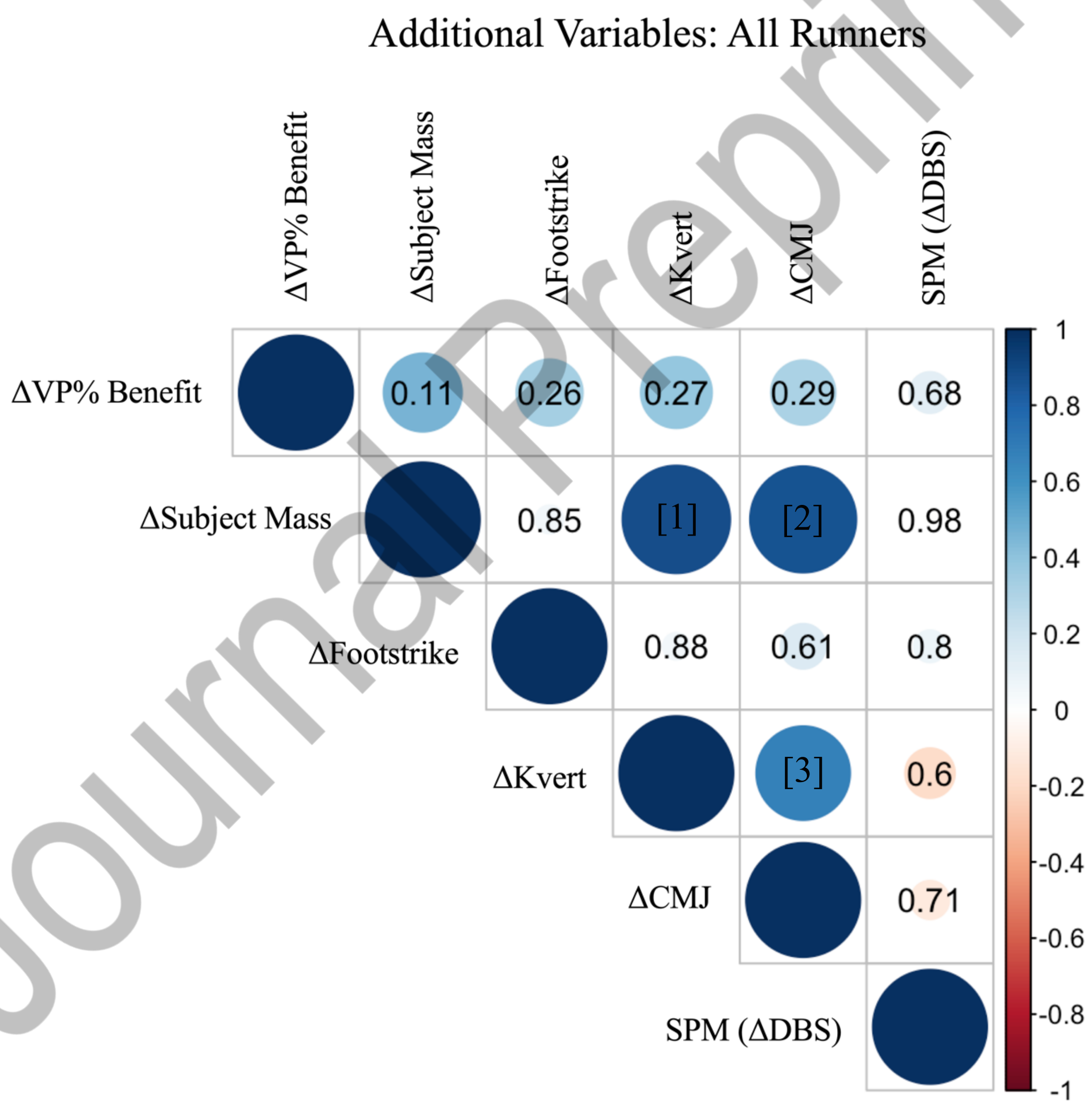


Journal Pre-proof

Figure S4. Correlation matrix showing the magnitude and direction of associations between changes in acute Vaporfly (VP) running economy improvements (ΔVP% benefit) and changes in biomechanical differences between shoes (ΔDBS), at the ankle, among VP trained runners from pre- to post-intervention. The magnitude and direction of each Pearson correlation (r) is illustrated by the colour and size of each circle, detailed by the vertical axis on the right. Non-significant p values are noted within each circle. Significant associations ($p < 0.05$), noted with a bracketed number, were: [1] p = 0.0165; [2] p = 0.0230; [3] p = 0.0435.

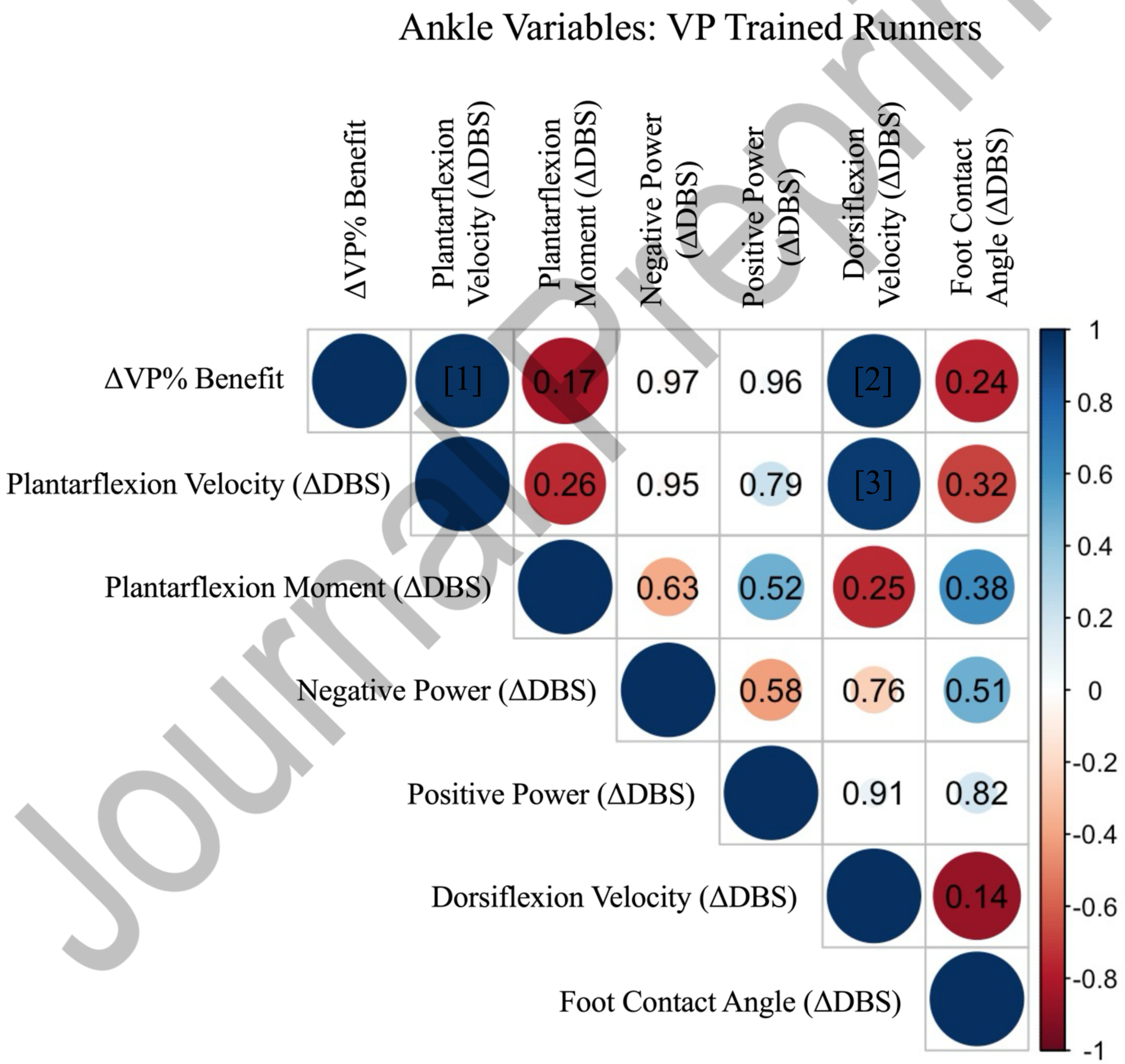


Journal Pre-proof

Figure S5. Correlation matrix showing the magnitude and direction of associations between changes in acute Vaporfly running economy improvements (ΔVP% benefit) and changes in biomechanical differences between shoes (ΔDBS), at the ankle, among traditional racing flat (FL) trained runners from pre- to post-intervention. The magnitude and direction of each Pearson correlation (r) is illustrated by the colour and size of each circle, detailed by the vertical axis on the right. Non-significant p values are noted within each circle. Significant associations ($p < 0.05$), noted with a bracketed number, were: [1] $p = 0.0451$; [2] $p = 0.0255$.

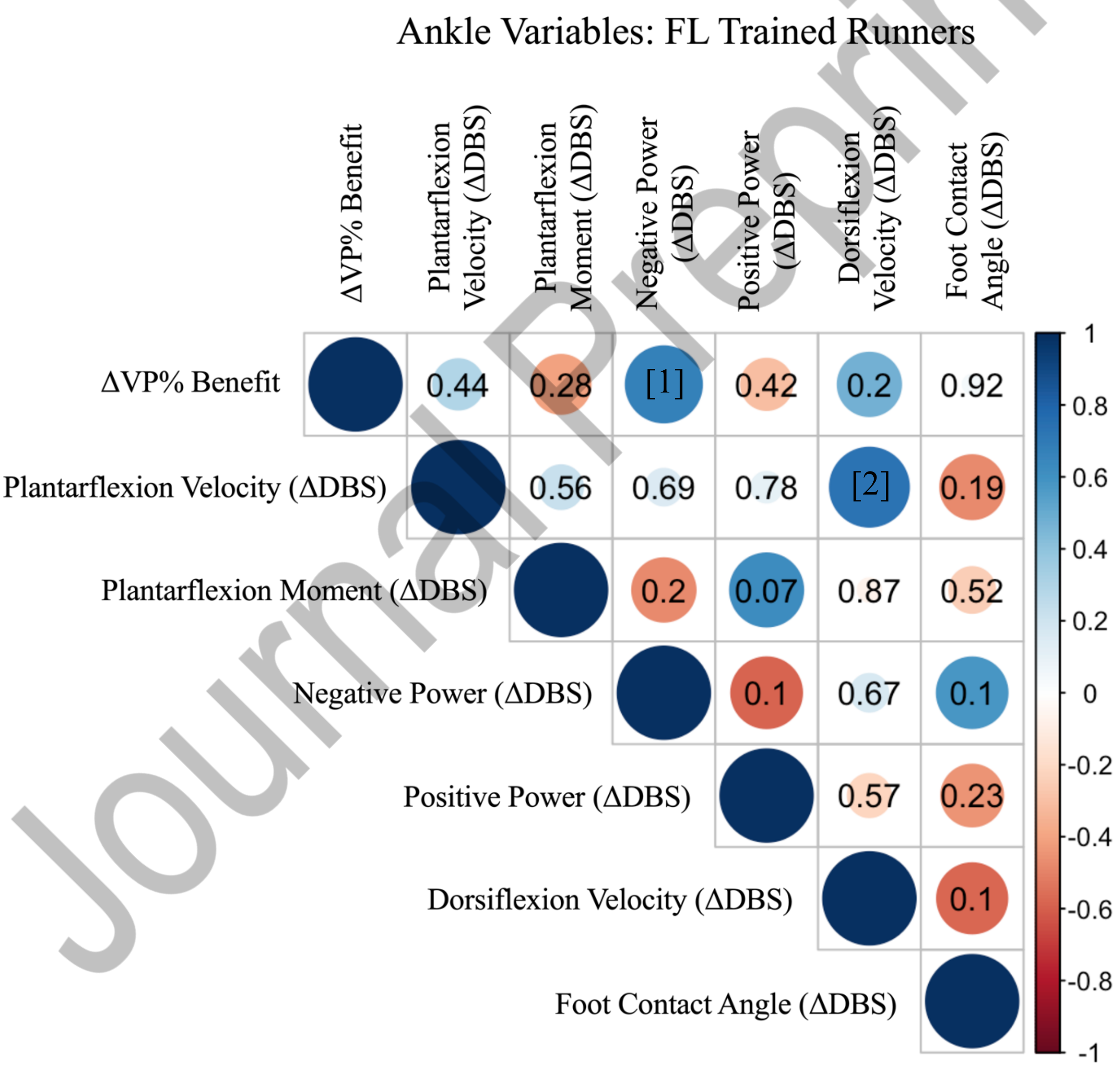


Journal Pre-proof

Figure S6. Correlation matrix showing the magnitude and direction of associations between changes in acute Vaporfly (VP) running economy improvements (ΔVP% benefit) and changes in biomechanical differences between shoes (ΔDBS), at the metatarsophalangeal (MTP) joint, among VP trained runners from pre- to post-intervention. The magnitude and direction of each Pearson correlation (r) is illustrated by the colour and size of each circle, detailed by the vertical axis on the right. Non-significant p values are noted within each circle. Significant associations ($p < 0.05$), noted with a bracketed number, were: [1] p = 0.0299; [2] p = 0.0408; [3] p = 0.0232; [4] p = 0.0347.

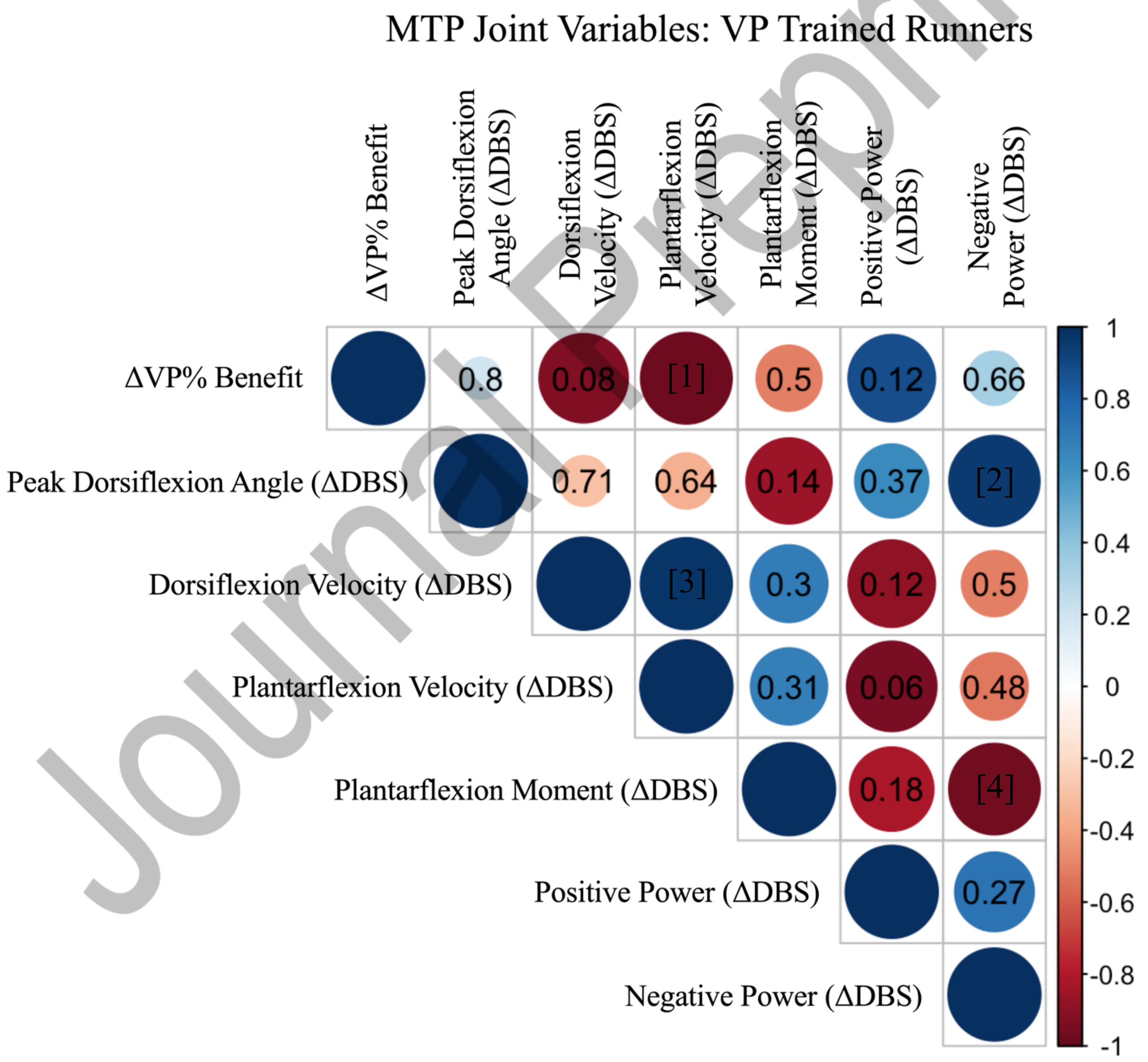


Journal Pre-print

Figure S7. Correlation matrix showing the magnitude and direction of associations between changes in acute Vaporfly running economy improvements (ΔVP% benefit) and changes in biomechanical differences between shoes (ΔDBS), at the metatarsophalangeal (MTP) joint, among traditional racing flat (FL) trained runners from pre- to post-intervention. The magnitude and direction of each Pearson correlation (r) is illustrated by the colour and size of each circle, detailed by the vertical axis on the right. Non-significant p values are noted within each circle. Significant associations ($p < 0.05$), noted with a bracketed number, were: [1] p = 0.0057; [2] p = 0.0271.

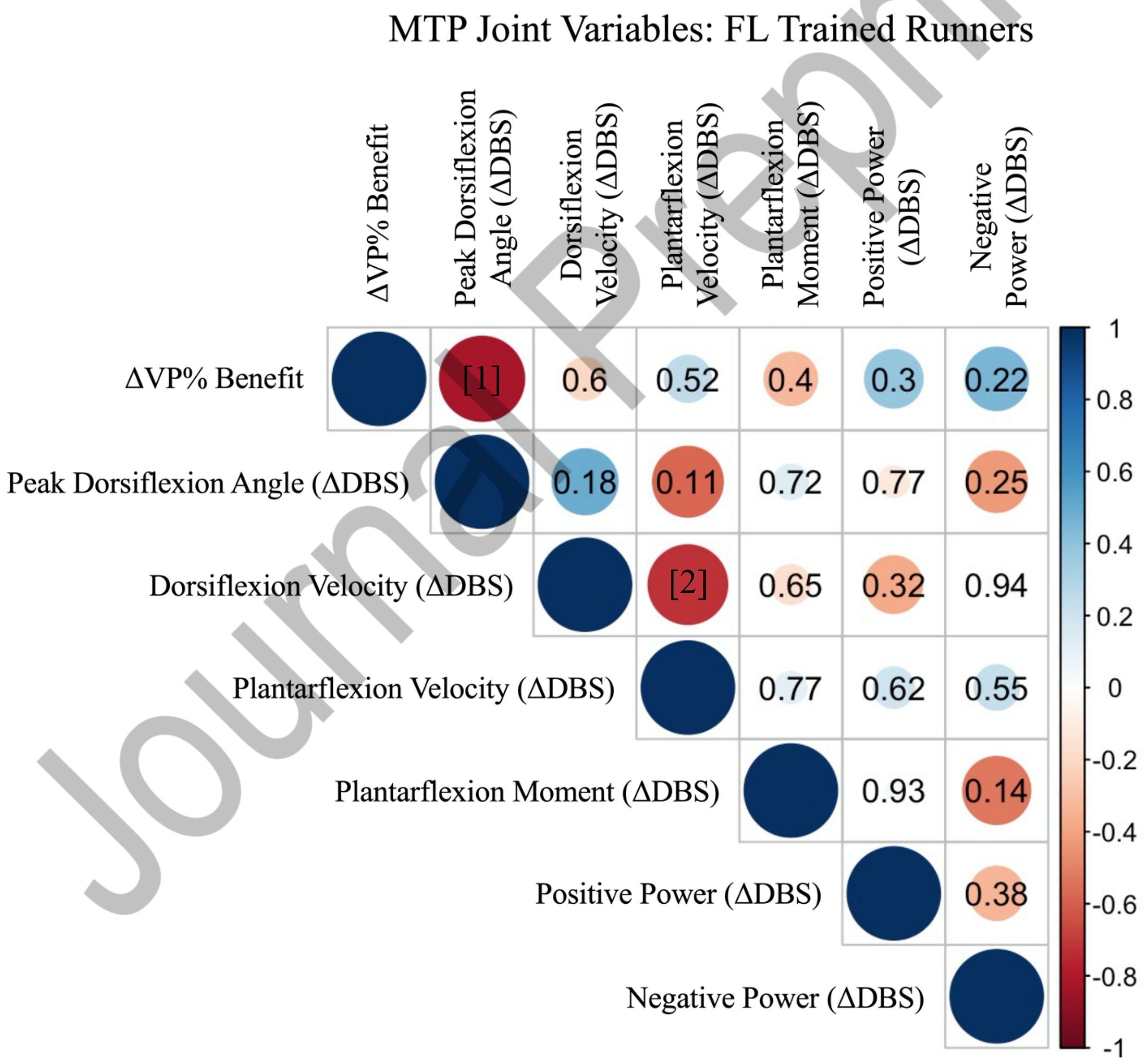


Journal Pre-print

Figure S8. Correlation matrix showing the magnitude and direction of associations between changes in acute Vaporfly (VP) running economy improvements (ΔVP% benefit) and additional variables of interest, among VP trained runners from pre- to post-intervention. The magnitude and direction of each Pearson correlation (r) is illustrated by the colour and size of each circle, detailed by the vertical axis on the right. Non-significant p values are noted within each circle. The significant association ($p < 0.05$), noted with a bracketed number, was: [1] $p = 0.0176$.

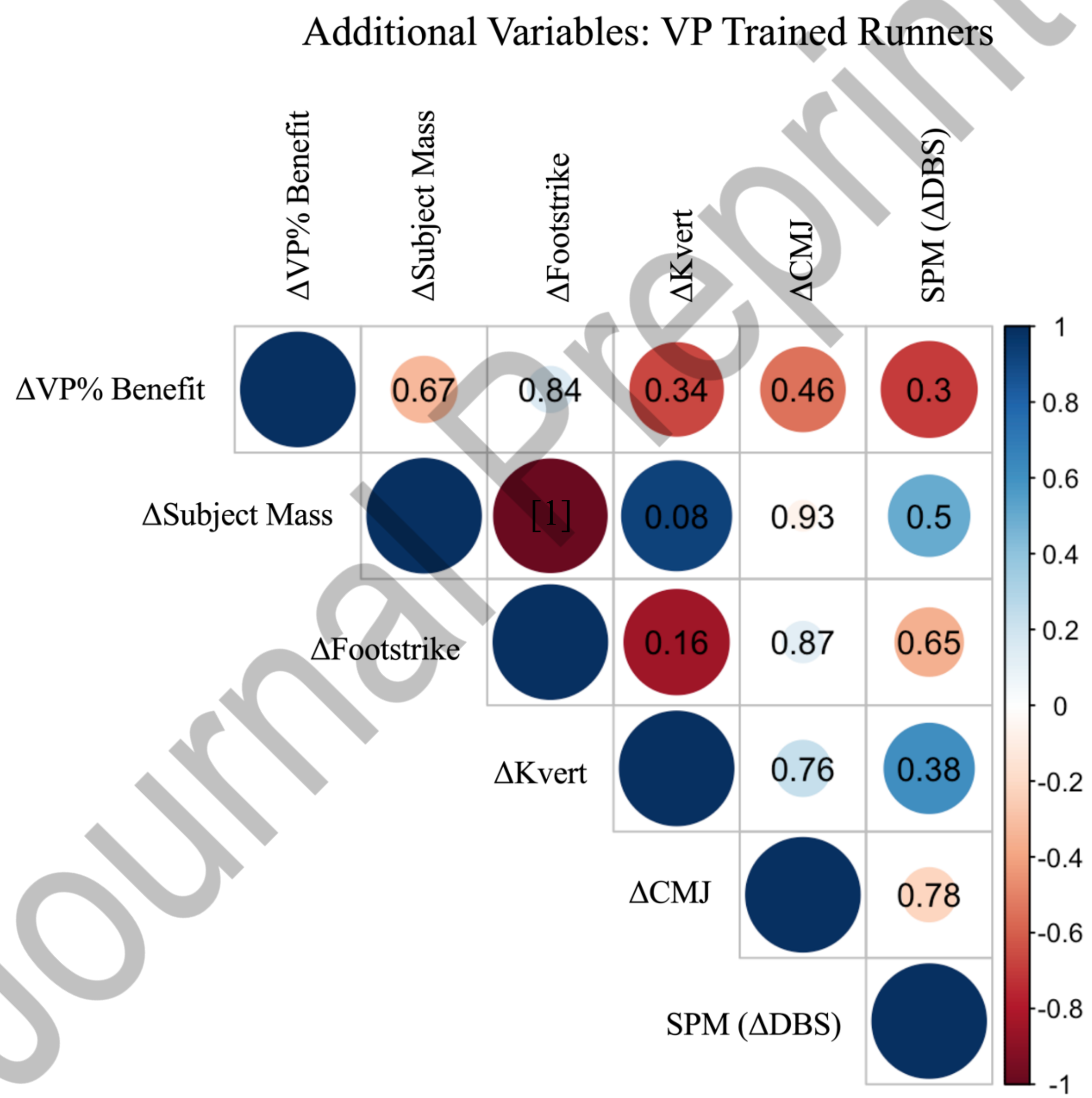


Journal Pre-proof

Figure S9. Correlation matrix showing the magnitude and direction of associations between changes in acute Vaporfly running economy improvements (ΔVP% benefit) and additional variables of interest, among traditional racing flat (FL) trained runners from pre- to post-intervention. The magnitude and direction of each Pearson correlation (r) is illustrated by the colour and size of each circle, detailed by the vertical axis on the right. Non-significant p values are noted within each circle. Significant associations ($p < 0.05$), noted with a bracketed number, were: [1] $p = 0.0138$; [2] $p = 0.0002$. Abbreviations: vertical hopping stiffness (Kvert), countermovement jump (CMJ), and steps per minute (SPM).

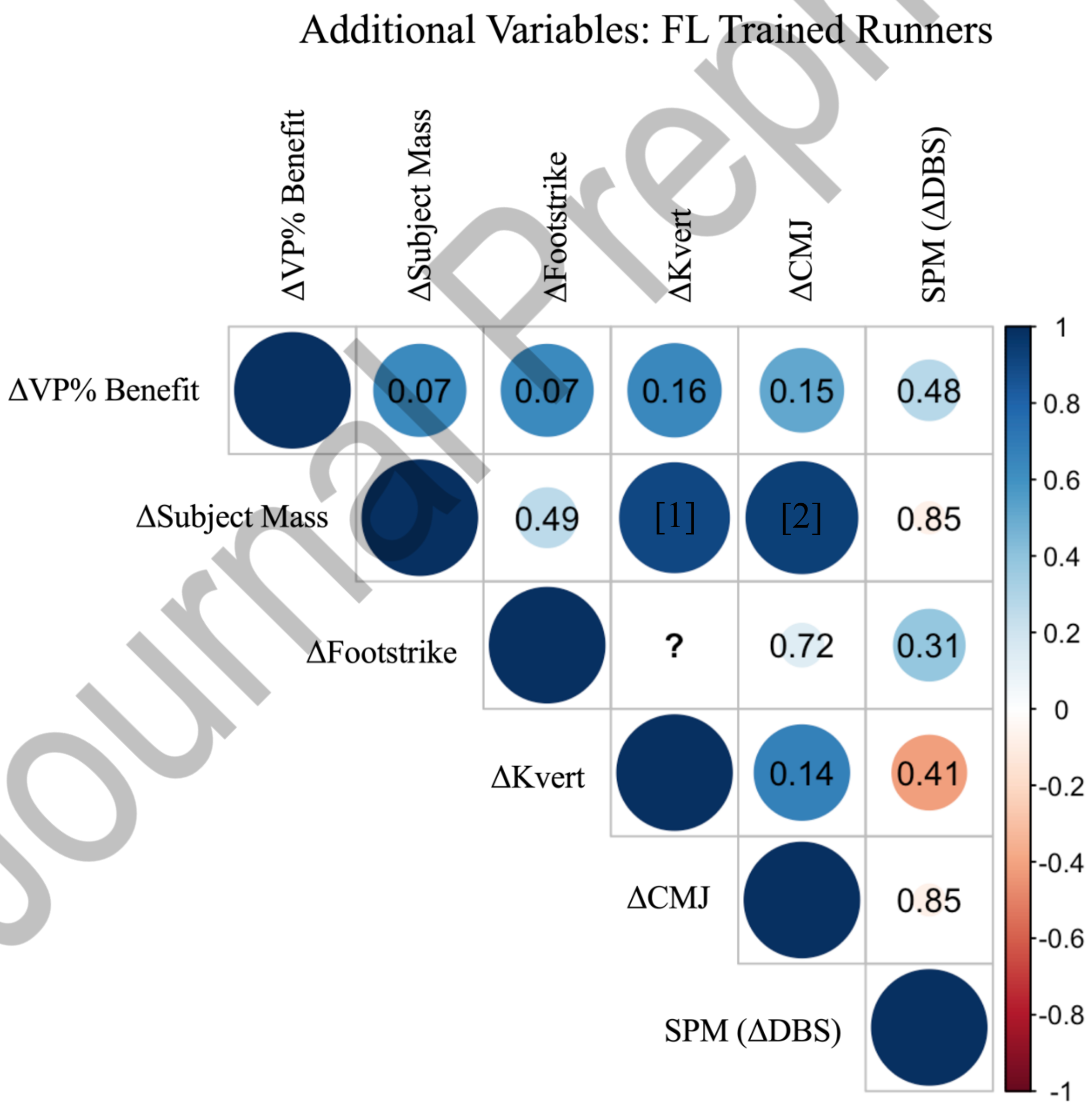


Journal Pre-proof
